# Response outcomes gate the impact of expectations on perceptual decisions

**DOI:** 10.1101/433409

**Authors:** Ainhoa Hermoso-Mendizabal, Alexandre Hyafil, Pavel E. Rueda-Orozco, Santiago Jaramillo, David Robbe, Jaime de la Rocha

**Affiliations:** IDIBAPS, Barcelona, Spain; Center for Brain and Cognition, Universitat Pompeu Fabra, Barcelona, Spain; Instituto de Neurobiología, UNAM, México; University of Oregon, Oregon,USA; Institut de Neurobiologie de la Méditerranée, Marseille, France

## Abstract

Perceptual decisions are not only determined by current sensory information but are also influenced by expectations based on recent experiences. Can the impact of these expectations be flexibly modulated based on the outcome of previous decisions? We trained rats in several two-alternative forced choice auditory tasks, where the probability to repeat the previous stimulus category was varied in blocks of trials. All rats capitalized on the serial correlations of the stimulus sequence by consistently exploiting a *transition bias:* a tendency to repeat or alternate their previous response using an internal trial-by-trial estimate of the sequence repeating probability. Surprisingly, this bias was null in trials immediately following an error. The internal estimate however was not reset and it became effective again causing a bias after the next correct response. This ability to rapidly activate and inactivate the bias was captured by a non-linear generative model of rat behavior, whereby a reward-driven modulatory signal gated the use of the latent estimate of the environment statistics on the current decision. These results demonstrate that, based on the outcome of previous choices, rats flexibly modulate how expectations influence their current decisions.

## INTRODUCTION

Imagine Rafa Nadal returning Roger Federer’s serve in the decisive game of a Grand Slam final. Serving at 185 km per hour, Nadal has a few hundred milliseconds to visually estimate the ball trajectory, prepare the motor plan including where he aims to return the ball and execute it. In such speeded decisions based on partial or ambiguous sensory information, the anticipation provided by an informed prior expectation can be decisive because subjects can respond faster. Based on past games bringing the two players together, and on the pattern of the last serves executed by Federer, Nadal inevitably forms an expectation about where the next ball will arrive. Combined with the visual motion of the ball, this expectation may allow him to gain some decisive tens of milliseconds in the return of the serve (Vernon et al., 2018). However, if his prediction fails and he concedes an ace, does he need to choose between trashing his prior model on Federer’s serve or sticking to it in the subsequent point? Or can Nadal transiently downplay the weight of his prediction on the next serve without modifying his prior?

Normative theories describe how prior expectations and ambiguous stimulus evidence should be combined in order to maximize categorization performance (Ernst and Banks, 2002; Stocker and Simoncelli, 2006). In dynamical environments, in which the statistics of the sensory information varies with time, subjects must be constantly updating their internal model by accumulating past stimuli, actions and outcomes (Yu and Cohen, 2008). The updating of the prior based on the actions occurring in each trial typically introduces sequential effects, which are systematic history-dependent choice biases reflecting the impact of the trial-to-trial variations in expectation (Abrahamyan et al., 2016; Akaishi et al., 2014; Akrami et al., 2018; Ashourian and Loewenstein, 2011; Braun et al., 2018; Busse et al., 2011; Cho et al., 2002; Fischer and Whitney, 2014; Fründ et al., 2014; Hwang et al., 2017; Meyniel et al., 2016; Nogueira et al., 2017). However, there are circumstances where subjects seem able to quickly and flexibly modulate the impact of prior expectations in driving their choices. One of such examples is the switch between (1) exploiting choices which, according to their current statistical model of the environment, are more likely to yield reward and (2) exploring alternative choices that are not aimed to maximize reward given that internal model, but to reduce environmental uncertainty and eventually refine the current model (Daw et al., 2006; Ebitz et al., 2018; Karlsson et al., 2012). In particular, when the task design potentiates stochastic exploration, rats are able to operate in an expectation-free mode in which choices did not depend on previous history (Tervo et al., 2014). In other tasks, the updating of the internal prior is not done in a continuous manner as new information is presented, but subjects update their internal estimates abruptly and intermittently when they feel there has been a change-point in the environment (Gallistel et al., 2004). Recent studies have shown that, in the absence of feedback, the magnitude of the expectation bias on current choice is smaller after a low confidence response (Braun et al., 2018; Samaha et al., 2018; Urai et al., 2017). Despite these findings, we lack a conceptual framework that could explain both how expectations are formed and which are the factors that regulate their use on a moment to moment basis.

Here we investigate whether the combination of expectation and sensory evidence can be dynamically modulated. Moreover, we aim to develop a unified model that jointly describes the dynamics of expectation build-up and its modulatory variables on a trial by trial basis. We trained rats to perform perceptual discrimination tasks using stimulus sequences with serial correlations. Behavioral analysis allowed us to tease apart the different types of history biases. In particular, rats accumulated evidence over previous choice transitions, defined as repetitions or alternations of two consecutive choices, in order to predict the next rewarded response. Crucially, this expectation-based bias disappeared after an error, reflecting a fast switch into an expectation-free categorization mode. This switch did not imply however the reset of the accumulated expectation which resumed its influence on behavior as soon as the animal obtained a new reward. This ubiquitous behavior across animals was readily captured by a non-linear dynamical model in which previous outcomes acted as a gate for the impact of past transitions on future choices.

## RESULTS

### A reaction time auditory discrimination task promoting serial biases

To study how the recent history of stimuli, responses and outcomes influence perceptual choices, we trained rats in a two-alternative forced choice (2AFC) auditory discrimination task in which serial correlations were introduced in stimulus trial sequences (Fig.1a-c) (Braun et al., 2018; Goldfarb et al., 2012; Jones et al., 2013; Kim et al., 2017). This design mimicked the temporal regularities of ecological environments and allowed us to probe the trial-by-trial expectations that animals formed about upcoming stimuli based on the changing statistics of the stimulus sequence. The serial correlations between trials were created using a two-state Markov chain (Fig. 1b) parameterized by the probability to repeat the previous stimulus category P_rep_ (the unconditioned probabilities for each of the two categories were equal). We varied P_rep_ between Repeating blocks, in which P_rep_= 0.7, and Alternating blocks in which P_rep_= 0.2 (Fig. 1c; block length 200 trials). By poking into the center port, rats triggered the presentation of the stimulus, which lasted until they freely withdrew from the port. Each acoustic stimulus was a superposition of a high-frequency and a low-frequency amplitude-modulated tones and animals were rewarded for correctly discriminating the tone with the higher average amplitude. The discrimination difficulty of each stimulus, termed stimulus strength *s*, was randomly and independently determined in each trial, and set the relative amplitude of each tone (Fig.1b,d). When stimulus strength *s* was null, i.e. contained no net evidence in favor of either alternative, the rewarded side was still determined by the outcome of the random Markov chain generating the stimulus category sequence (Fig. 1b).

**Figure 1.**
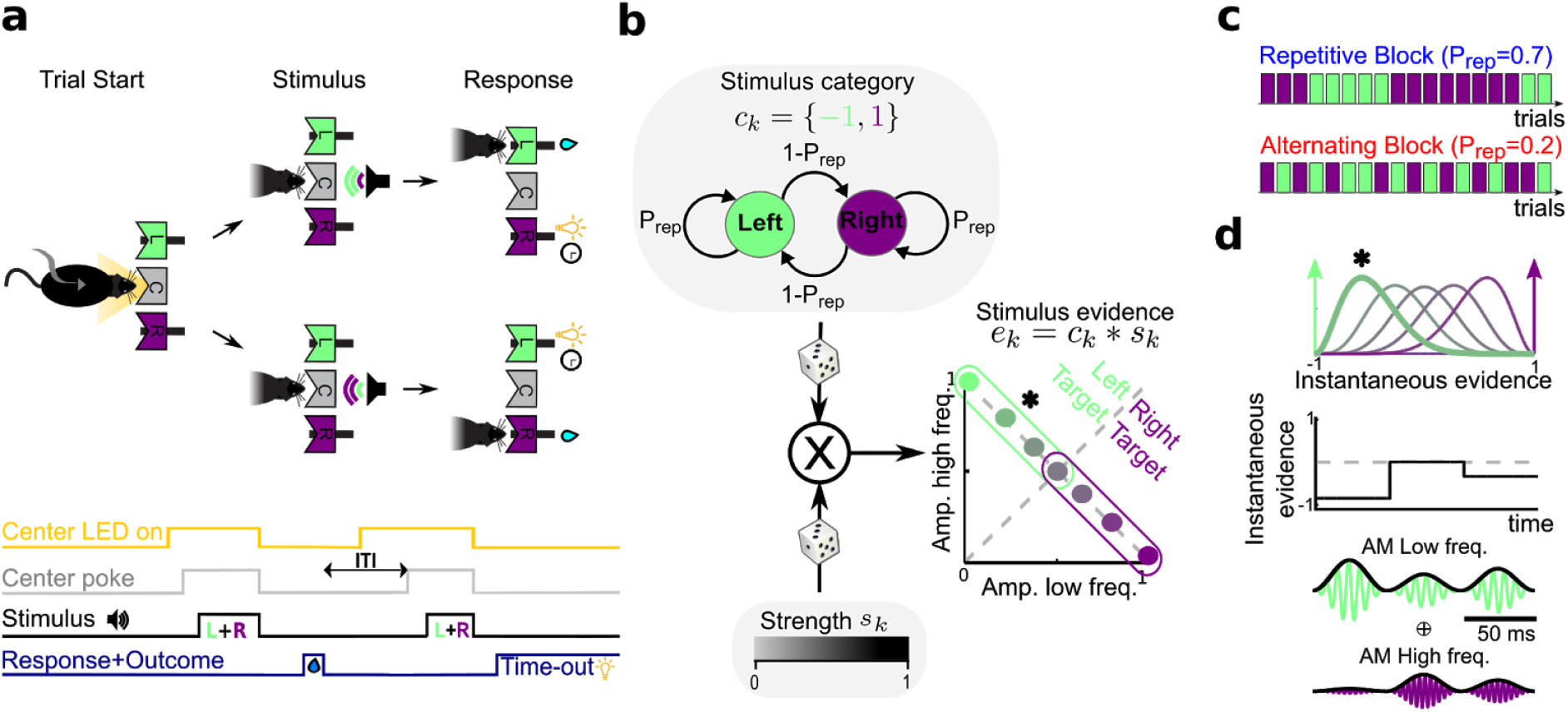
Auditory discrimination task and stimulus sequence statistics. **a**, Sketch of one trial of the task: cued by center port LED, rats poke in the center port to trigger the presentation of a mixture of two AM tones, each of which is associated with reward in the Left (L) or Right (R) port. Correct responses are rewarded with water and incorrect responses are punished with a light plus a 5 s time-out. **b-c**, Serial correlations in the sequence of stimuli were introduced by setting the probability of repeating the previous stimulus category *P*_*rep*_ (top in b) in blocks of 200 trials named Repetitive Block and Alternating Block (c). The stimulus strength *s*_*k*_ was randomly drawn in each trial (bottom in b) to yield the stimulus evidence *e*_*k*_, that determined the distance to the categorization boundary, i.e. the discrimination difficulty of the stimulus (right in b). **d**, The stimulus evidence *e*_*k*_ determined the distribution (top) from which the instantaneous evidence was drawn in each frame of the sound envelope (see color match with b). An instantaneous evidence trace (middle) and the AM modulated tones that result (bottom) are shown for an example stimulus with *e* = −0.48 (asterisks in b and d).

### Across-trial dynamics of history-dependent choice biases

Animals in Group 1 (n=10 animals) completed an average of 508 trials per session (range 284 - 772 average trials), gathering an average of 56,242 trials in total per animal (range 15,911 - 81,654 trials). Psychometric curves showing the proportion of Rightward responses as a function of the stimulus evidence did not depend on block type (Fig. 2a left). To estimate the impact of stimulus serial correlations, we also analyzed the repeating psychometric curves, showing the proportion of trials where the animals repeated the previous choice as a function of the sensory evidence in favor of the repeating choice. This analysis showed that all animals developed a block-dependent repeating bias *b* (Fig. 2a right, b left): after correct trials, *b* was positive in the Repetitive Block, and negative in the Alternating Block (i.e. tendency to repeat or alternate, respectively). Interestingly, the fixed side bias *B*, measuring the history-independent preference to choose one side, was similar across blocks for each animal (Fig. 2c left), showing that animals side preference was independent of the changes in repeating bias caused by block switching. Surprisingly, in trials following an error, *b* almost vanished in both block types (Fig. 2b-c right). Thus, after errors rats did not use previous history to guide their decision (e.g. in the Repetitive block, after an incorrect Rightward response, the Leftward response is more likely to be rewarded). In contrast, the fixed side bias *B* was analogous after correct and error trials (Supplementary Fig. 1b).

**Figure 2.**
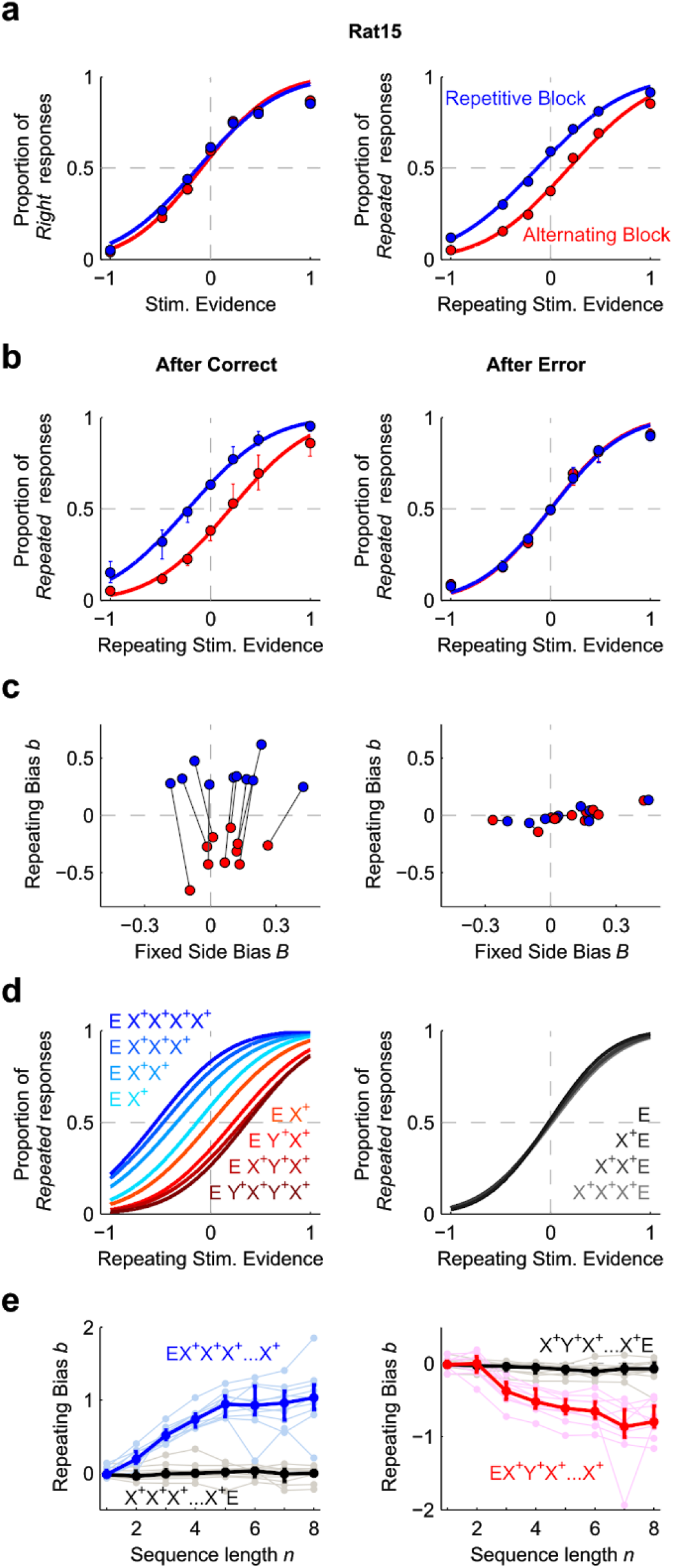
Build-up and reset dynamics of the repeating bias. **a**, Psychometric curves for an example animal showing the proportion of Rightward responses vs. stimulus evidence (left) or of Repeated responses vs. Repeating stimulus evidence (right) computed in the Repetitive (blue dots) or Alternating blocks (red dots; color code applies for all panels). This animal shows a block independent Rightward fixed side bias *B*>0 (left), and a block-dependent repeating bias *b* matching the tendency of each block (right). Curves show fits using a probit function. **b**, Proportion of Repeated responses (median across *n* = 10 animals) computed in trials following a correct (left) or an incorrect response (right). **c**, Repeating bias *b* versus fixed side bias *B* in the two blocks after a correct (left) or an incorrect response (right). Each pair of connected dots represents one animal. **d**, *Left:* Fits of the proportion of Repeated responses following trial sequences made of a different number of correct repetitions (blue gradient) or alternations (red gradient; see insets for color code). X^*+*^ and Y^*+*^ represent either Rightward or Leftward correct responses. E represents an error. Time in the sequences progresses from left to right. *Right:* same curves obtained when the sequence of correct repetitions is terminated by an error. **e**, Repeating bias versus the sequence length of correct repetitions (left, blue) or alternations (right, red). Sequences terminated by an error are shown in black. Dark traces show median across animals while light traces show individual animals. Error bars show Std. Dev. (a) or first and third quartiles (b, e).

Rats used history information by tracking several trials over short windows into the past: the magnitude of the repeating bias *b* built up with the number of consecutive correct past repetitions or alternations *n* until it plateaued after *n =* 5-10 trials (Fig. 2e blue and red line). This plateau was greater after repetitive patterns rather than alternating patterns. Importantly however, irrespective of *n*, the repeating bias *b* reset almost completely with a single incorrect response for all rats (Fig. 2e). The reset occurred independently of the strength of the incorrectly categorized stimulus (Supplementary Fig. 1c). To control that the reset was not caused by forgetting due to the 5s time-out imposed after errors, we trained a subset of rats using random time-out durations (range 1-5 s) and found that the bias reset was maintained independently of time-out duration (Supplementary Fig. 1d). The reset was only observed after errors and not after correct but unexpected responses, e.g. one alternation after several repetitions (Supplementary Fig. 2a). Accordingly, performance was higher for trials following a correct trial than for trials following an error, and it increased with *n* (Supplementary Fig. 3a-b). Moreover, the repeating bias increased performance when it was consistent with the block tendency but it decreased performance when it was inconsistent with it (Supplementary Fig.3c-d). The impact on performance was largest for low stimulus strength, when the sensory evidence was weak and animals relied more strongly in their *a priori* belief, and it vanished to zero as the stimulus strength increased and the classification became easier. Together, these observations suggest that rats update their beliefs about the environment in a trial-by-trial basis and that this update crucially relies on the outcome of the preceding trials: longer sequences of rewarded repetitions/alternations lead to stronger response prior, but one error was sufficient to make the animals abandon this prior.

### A GLM analysis of integration of sensory evidence and recent history information

Having identified that rats used previous responses and outcomes to guide decisions, we aimed to identify the specific factors in trial history generating this repeating choice bias. In particular, these factors could be (1) a *lateral bias* that creates an attraction or repulsion towards the Left or Right side depending on previous responses (Fig. 3a) and (2) a *transition bias* that creates an attraction towards repeating or alternating depending on the history of previous repetitions and alternations (Fig. 3c). To understand the difference between these first-order (lateral) and second-order (transition) biases, we first considered correct responses only, and described the effect of errors below. If subjects were using e.g. the last four choices to estimate the probability of the next stimulus category, given the example choice sequence *R*^*+*^*R*^*+*^*R*^*+*^*L*^*+*^, where *R*^*+*^ and *L*^*+*^ represent a Rightward or Leftward correct choice (*L*^*+*^ represents the last trial, Fig. 3b), they would estimate that R is more likely and develop a lateral Rightward bias γ^*L*^ in the next trial (Fig. 3a). The same four-choice sequence can however be represented as the series of transitions *Rep*^++^*Rep*^++^*Alt*^++^, where *Rep*^++^ and *Alt*^++^ represent repetitions and alternations between two correct responses. These transitions sequence is first accumulated into the *transition evidence z*^*T*^, an internal estimate of the probability of the next transition, which in this example points the subject to predict a Repetition in the last trial (Fig. 3c). Importantly, the transition evidence *z*^*T*^ needs to be converted into an effective decision bias by projecting it into the Right-Left choice space (Fig. 3c-d). This is achieved by multiplying *z*^*T*^ with the last response *r*_*t-1*_, yielding the *transition bias γ*^*T*^ = *z*^*T*^ × *r*_*t*-1_ (see gray arrows in Fig. 3b-d). Lateral and transition biases have an opposite influence in the final choice: while γ^*L*^ has a Rightward influence, γ^*T*^ has a Leftward influence because the transition evidence *z*^*T*^ predicts a repetition and the last choice was Leftward (compare Fig. 3a and d). Thus, the two biases extract different information from the sequence of past trials. Although only the transition bias is adaptive in the task, since it allows to capitalize the sequence correlations in both types of blocks, the two biases could in principle contribute to the repeating bias *b* described above.

**Figure 3.**
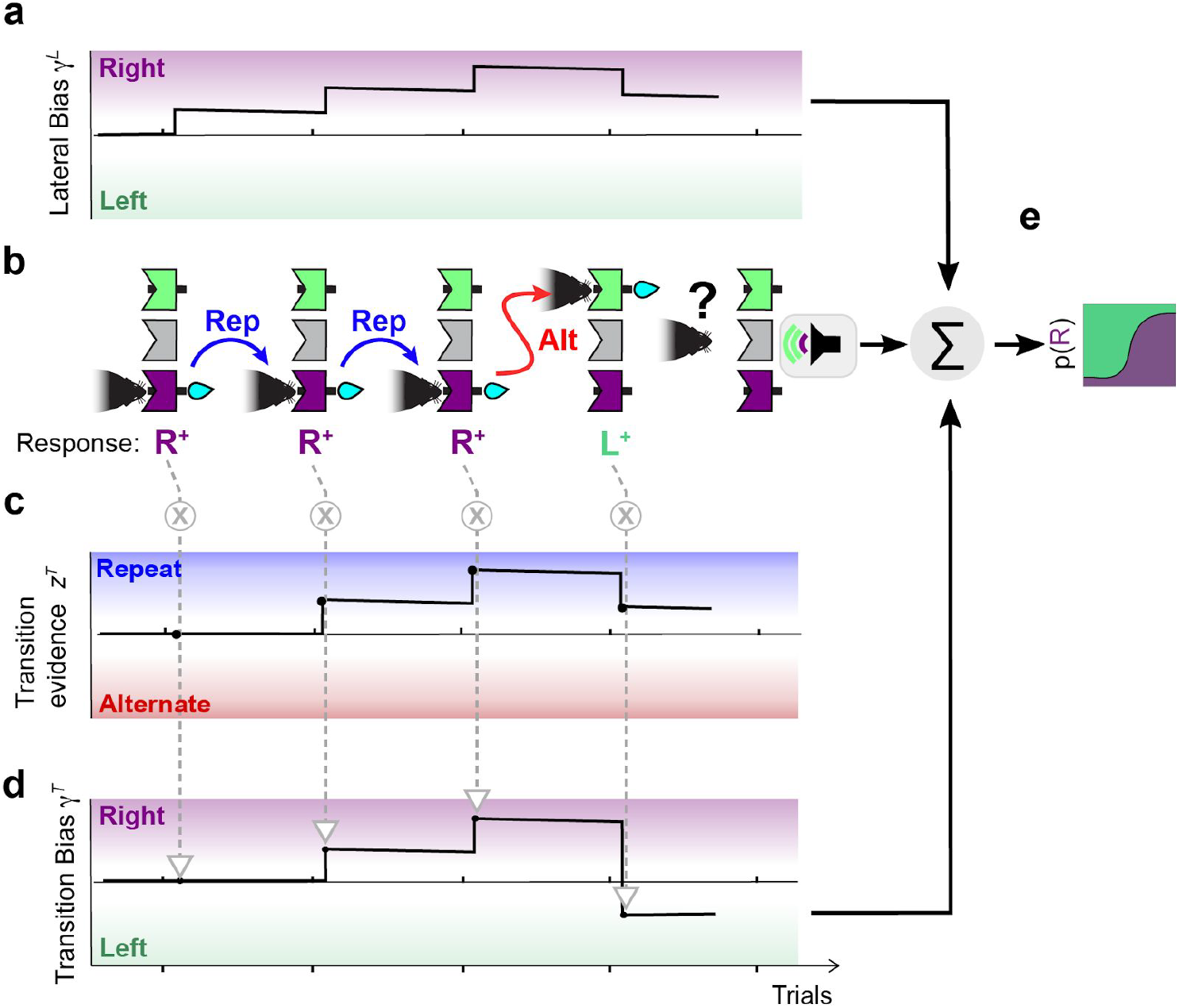
Dissecting two different history choice biases. Cartoon of an example series of four choices, *R*^*+*^*R*^*+*^*R*^*+*^*L*^*+*^, illustrating the build-up of the lateral and transition biases. **a**, The lateral bias, capturing the tendency to reinforce Rightward or Leftward rewarded responses, increases towards the Right in the first three *R*^*+*^ trials and compensates this build-up with the last *L*^*+*^ response. Its net impact on the final trial is a Rightward bias. **b**, Schematic of the sequence of rewarded rat responses showing the transitions, defined as the relation between two consecutive responses, being Repetitions (Rep, blue arrows) or Alternations (Alt, red arrow). For each choice, the animal combines its expectation based on previous trials with current stimulus sensory information (see last trial). **c**, Transition evidence *z*^*T*^ accumulates the series of transitions *Rep*^++^*Rep*^++^*Alt*^++^ predicting a Repetition in the final trial. **d**, The transition bias *γ*^*T*^ is obtained by projecting the Transition evidence *z*^*T*^ (c) onto the Right-Left choice space (see gray arrows). **e**, The evidence provided by the current stimulus is summed to the addition of the biases *γ*^*L*^_*t*_ + *γ*^*T*^_*t*_ and passed through a sigmoid function, yielding the probability of selecting a Rightward response (Supplementary Fig. 4).

To quantify the impact onto current decisions of the lateral and the transitions biases, and to investigate their dependence on error responses, we used a generalized linear model (GLM) (Abrahamyan et al., 2016; Braun et al., 2018; Busse et al., 2011; Fründ et al., 2014; Nogueira et al., 2017; Urai et al., 2017). The GLM separately measured the impact onto the current decision of each response *r* (*r = R, L*) and transition *T* (*T = Rep, Alt*) in the last ten trials (Supplementary Fig. 4; see Methods for details). Because correct and error choices presumably had a different impact (Fig. 2b-d right), we separated the contribution to the lateral bias γ^*L*^ of rewarded responses *r*^*+*^, sometimes called *reinforcers (Corrado et al., 2005; Lau and Glimcher, 2005)*, from error responses *r*^*−*^ (Supplementary Fig. 4). Following the same rationale, we separated the contributions to the transition bias γ^*T*^ of two consecutive correct responses (*T*^++^) from transitions where either the first (*T*^*−+*^), the second (*T*^+−^) or both responses (*T*^*−−*^) were incorrect. After fitting the regression weights of the GLM individually for each rat, a consistent pattern across animals emerged (Fig. 4 orange curves and Supplementary Fig. 5a-d). The contribution of each response to γ^*L*^ depended on its outcome following a win-stay-lose-switch pattern: while rats displayed a tendency to opt again for the side of previously rewarded responses (positive *r*^*+*^ weights), they tended to opt away from previously non-rewarded responses (negative *r*^*−*^ weights; Fig. 4a orange curves). Similarly, previous transitions between two correct responses *T*^++^ were positively weighted (Fig. 4b-c orange curves), meaning that recent ++ repetitions increased the tendency to repeat (positive impact on γ^*T*^), while recent ++ alternations increased the tendency to alternate (negative impact on γ^T^). However, the transitions *T*^+−^, *T*^*−+*^ and *T*^*−−*^ with at least one error barely influenced subsequent choices (Fig. 4b). This means that, in the example choice sequence *R*^*+*^*R*^*+*^*R*^*+*^*R*^*+*^, equivalent to the transition sequence *Rep*^++^*Rep*^++^*Rep*^++^, only the first two repetitions impacted on γ^*T*^. Thus, the only effective transitions driving the transition bias were ++ transitions.

**Figure 4.**
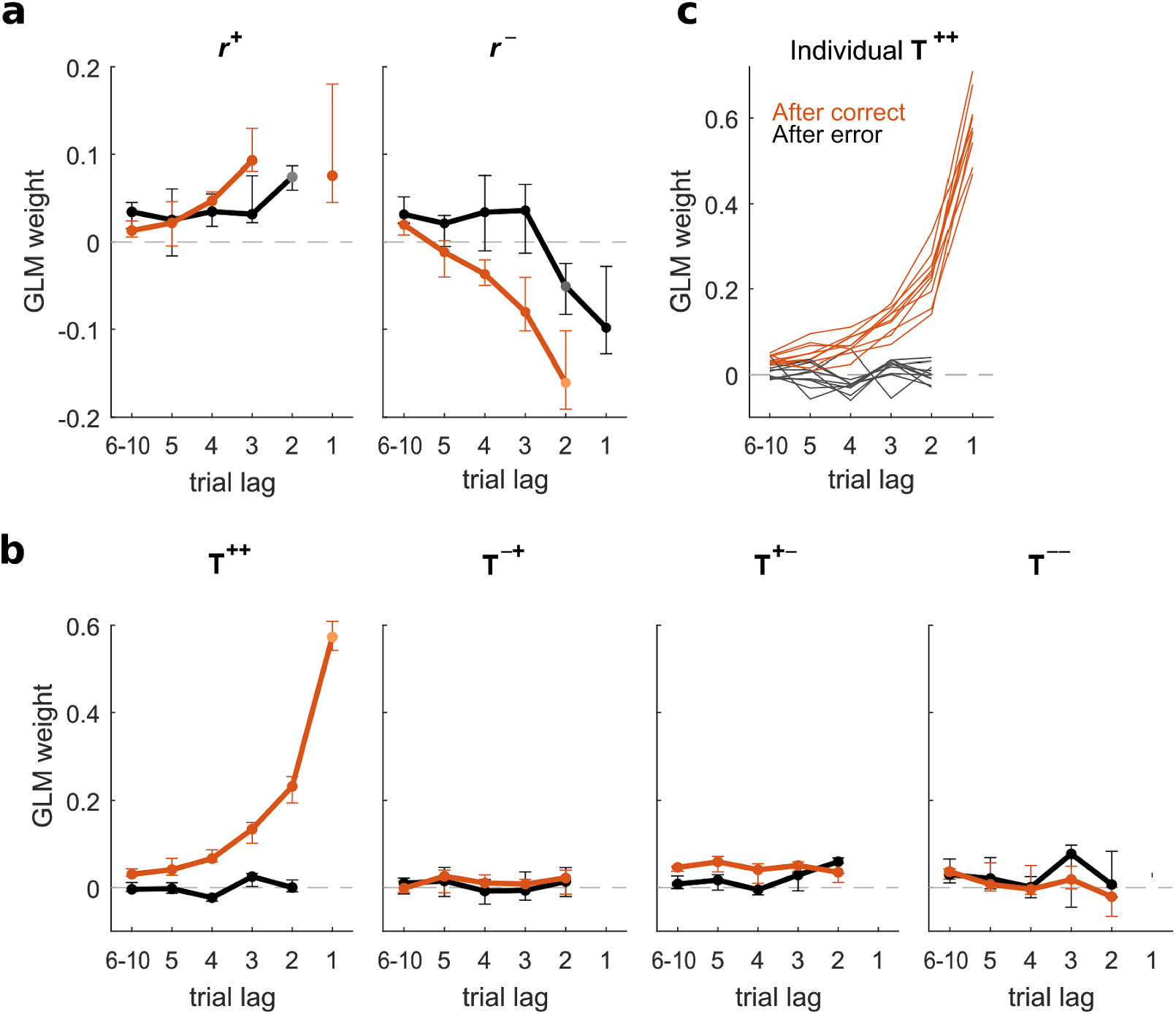
Fitted weights quantifying the impact of the lateral and transition biases onto animals decisions. Coefficients obtained in the GLM when separately fitting the choices in trials after a correct (orange) and error response (black). **a**, Influence of the response side (Left vs Right) from previously rewarded (*r*^*+*^, left panel) and unrewarded (*r*^*−*^, right panel) trials. **b**, Influence of previous transitions (repetition vs. alternation) computed separately for *T*^++^(a rewarded trial followed by a rewarded trial), *T*^*−+*^ (error-rewarded), *T*^+−^(rewarded-error) and *T*^*−−*^ (error-error). Points in a-b show median coefficients across animals (*n* = 10) and error bars indicate first and third quartiles. **c**, Transition kernels for individual animals show the ubiquity across subjects of the reset of the kernel after errors.

Error responses had yet a more dramatic effect on the transitions bias. They not only made the *T*^+−^, *T*^*−+*^ and *T*^*−−*^ transitions ineffective but they also suppressed the impact of all previously accumulated *T*^++^ transitions: the weights of previous *T*^++^ transitions were completely vanished when we fitted the GLM only using choices following an error trial (Fig. 4b-c, black curves). Thus, after an error choice, the transition bias was reset to zero, γ^*T*^ = 0, meaning that rats behavior was completely blind to the history of previous repetitions and alternations, and was driven only by sensory information and lateral bias. The reset of γ^*T*^ was not an idiosyncratic strategy followed by some of our animals but it was found in every animal we trained (Fig. 4c and Supplementary Fig. 6). In fact the magnitude of transition kernel was much more homogenous across animals than the lateral kernel (Supplementary Fig. 6). The reset effect was however not observed in the lateral bias, which was moderately affected by errors (Fig. 4a, black curves). Thus, the bias reset following errors was specific to the transition term and extremely reliable across subjects.

Despite the strong impact of the transition bias, animal choices mostly relied on the current stimulus, which had an impact an order of magnitude larger than the transition bias, which was itself an order of magnitude larger than lateral bias (Supplementary Fig. 5e). The weakest (yet very consistent) sequential component was a stimulus repulsive bias reminiscent of an after-effect caused by sensory adaptation with a very slow recovery (Supplementary Fig. 5b). A modified analysis separating the effects of repetitions and alternations showed that they had largely symmetrical effects, suggesting that animals summarized both types of transition into a single rule that could take positive or negative values (Supplementary Fig. 7c). Importantly, the weights were identical when computed separately in repetition and alternation blocks (Supplementary Fig. 8). This suggests that rats adopted a single strategy across all blocks, and the different repeating choice bias found in each block (Fig. 2b-e) simply reflected the difference in the statistics of the stimulus sequence (Fig. 1c). Because the impact of transitions decayed in around 5 trials (Fig. 4b left), the strategy allowed animals to switch the sign of their repeating bias relatively fast when switching between blocks (Supplementary Fig. 1a) at the cost of suffering relatively large fluctuations in the repeating bias within each block. Model comparisons further confirmed that the full model fitted separately for trials following correct trials and errors provided a better fit to rats decisions than the full model fitted to all trials, or alternative models where the lateral and/or transition module were removed (Supplementary Fig. 7a). Importantly, the GLM with only lateral biases yielded an non-monotonic kernel for the Lateral responses, a result that could lead to spurious interpretations when the effect of previous transitions was not considered (Supplementary Fig. 7b).

To test the extent to which these findings depended on the task design, we trained a new group of rats (Group 2, *n* = 6) in a different level discrimination 2AFC task in which noise stimuli had to be classified according to the intensity difference between the two lateral speakers (Pardo-Vazquez et al., 2018). The stimulus sequence followed the same pattern as before with repeating and alternating blocks (Fig. 1b-c). Performing the same GLM analysis in this task yielded qualitatively the same results, including the reset of the transition bias after errors (Supplementary Fig. 9). Finally, we found that the presence of a transition bias and its reset after errors was not contingent on the presence of serial correlations in the stimulus sequence. A third group of rats (Group 3, *n* = 9) exposed to only an uncorrelated stimulus sequence, exhibited the same qualitative pattern for the impact of previous transitions, although of smaller magnitude (Supplementary Fig. 10c). Once the sessions started featuring stimulus serial correlations (Fig. 1b-c), the magnitude of the transition weights increased (Supplementary Fig. 10c) suggesting that the transition bias is an intrinsic behavior of the animals, but its magnitude can be adapted to the statistics of the environment. In total, these analyses show that the dependence on previous outcome of history-dependent biases is a general result across animals and across different tasks.

### Transition evidence is blocked but not reset after an error

We then asked whether the reset of the transition bias after errors reflected (i) a reset of the transition accumulated evidence *z*^*T*^, meaning the entire system monitoring transitions underwent a *complete reset* (Fig. 5a); or whether, in contrast, (ii) information about previous transitions was maintained in *z*^*T*^ but was gated off from causing a transition bias (Fig. 5b). Whereas in the latter scenario (*gating hypothesis)*, the information maintained in *z*^*T*^ could be used to compute γ^T^ following new correct responses, in the complete reset scenario the build-up of both *z*^*T*^ and γ^T^ started back from zero following errors. To test these two hypotheses, we quantified how the value of the bias γ^T^ in trial *t* could predict the bias at trial *t*+1, *t+2* and further, depending on the outcome of each of these trials. When trial *t* was correct, the bias was passed on to *t+1* with a discounting decay that mirrored the shape of the transition kernel in the GLM analysis (Fig. 5c dark orange dots). The same discounting occured going from *t*+1 to *t*+2 when trial *t*+1 was correct. By contrast, if *t* was incorrect, because of the bias reset after errors, the value of γ^T^ was not predictive of the decision at trial *t+1*, nor at trial *t+2* if *t+1* was also incorrect (Fig. 5c black dots). Crucially though, the bias γ^T^ in trial *t* strongly influenced choices at trial *t+2* if trial *t* was incorrect but trial *t+1* was correct (Fig. 5c light orange dots). Its impact was significantly larger than zero for all rats (Wald test *p* < 0.003 for each of the *n* = 10; Supplementary Fig. 5g) and close in magnitude to the impact when both trials *t* and *t+1* were correct. This rebound in choice predictability was even observed at *t+*3 after two incorrect responses followed by a correct one (Wald test *p* < 0.05 for nine out of the *n* = 10). These results are consistent with the gating hypothesis (Fig. 5b) in which errors do not cause a reset of the accumulated transition evidence *z*^*T*^ but do cause a transient cut off in the influence of *z*^*T*^ on choice, visible as a reset in γ^T^. This influence became effective again once the animal made a new correct response giving rise to the measured correlation between the values of the bias before and after the reset (Fig. 5b gray vertical arrows; Supplementary Fig. 5h). An equivalent analysis on the lateral bias γ^L^ showed that the bias transferred to the subsequent trials with a rapid decay, which was moderately affected by the outcome of the trials and showed no evidence of reset-and-rebound dynamics (Supplementary Fig. 5f).

**Figure 5.**
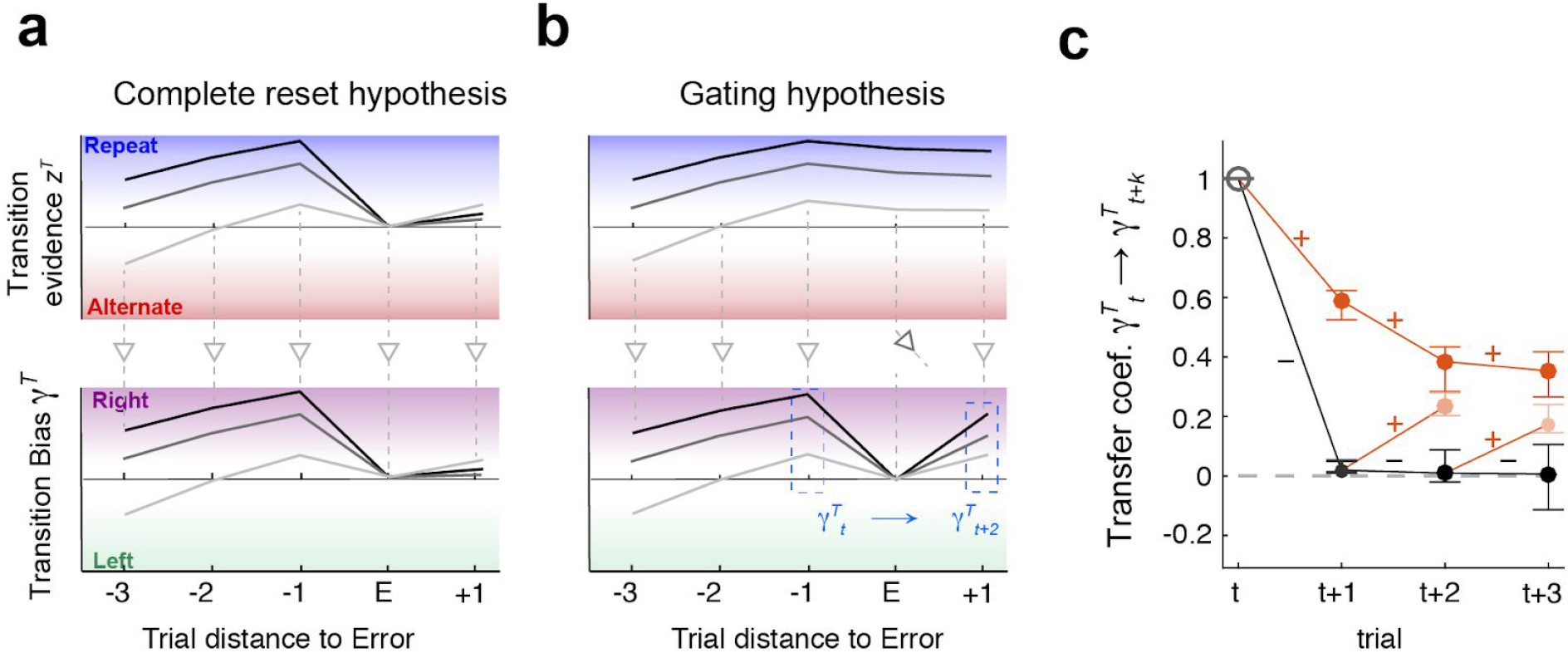
Transition bias is reset after errors but accumulated transition evidence is maintained. **a-b**, Schematics showing three example traces of the transition accumulated evidence *z*^*T*^_*t*_ (top) and transition bias on current response γ^*T*^_*t*_ (bottom) in two hypothetical scenarios. **a**, Complete reset hypothesis: after an error at *t*, both variables reset *z*^*T*^_*t*+1_ ≃ *γ*^*T*^_*t*+1_ ≃ 0. Evidence *z*^*T*^_*t*+2_ is then built up *de novo*, implying that biases before (γ^*T*^_*t*_) and after (γ^*T*^_*t*+2_) the reset are uncorrelated (γ^*T*^_*t*_ traces do not maintain the sorting). **b**, Gating hypothesis: after an error, evidence *z*^*T*^_*t*+1_ is maintained but it does not convert into a bias, leading to the reset γ^*T*^_*t*+1_ ≃ 0. After a correct response at *t*+1 the conversion is recovered and the value γ^*T*^_*t*+2_ correlates with γ^*T*^_*t*_ (γ^*T*^_*t*_ traces maintain the sorting). **c**, Transfer coefficient γ^T^_t_ → γ^T^_t+k_ versus trial lag *k* quantifies the degree to which the transition bias at trial *t* correlates with the bias on subsequent trials (blue dashed boxes in b). It is calculated separately depending on the outcome of each trial (colored lines show rewarded choices and black lines error choices; see Supplementary Methods for details). While the transfer coefficient vanishes after errors (i.e. reset of the bias; black dots), a correct response following an error (light orange) brings it close to the value obtained when there are no errors (dark orange dots). This implies that the information about the value of the bias γ^*T*^_*t*_ is maintained when the bias is reset (i.e. gating hypothesis).

### A dynamical model of history-dependent outcome-gated biases

Having found that the transition bias underwent reset-and-rebound dynamics, we built a generative model that could implement the gating hypothesis. One latent variable in the model was the accumulated transition evidence *z*^*T*^, which was updated in each trial depending on whether the last choice was a repetition or an alternation and therefore maintained a running estimate of the transition statistics (Braun et al., 2018; Busse et al., 2011; Cho et al., 2002; Fründ et al., 2014) (Fig. 6a). The dependence of the leak of *z*^*T*^ on the previous outcome could in principle implement the *Complete reset* hypothesis (Fig. 5a) if the leak following errors was complete (λ_T_ ≃ 1). A second modulatory variable *c*^*T*^ modulated the influence of the transition evidence onto the current decision by setting the transition bias equal to γ^*T*^ = *c*^*T*^ × *z*^*T*^ × *r*_*t*−1_. Importantly *c*^*T*^ was updated after each trial based on the trial outcome. In addition to the transition bias, the model also featured accumulated lateral evidence *z*^*L*^ that directly resulted in a lateral bias (i.e. γ^*L*^ = *z*^*L*^).

**Figure 6.**
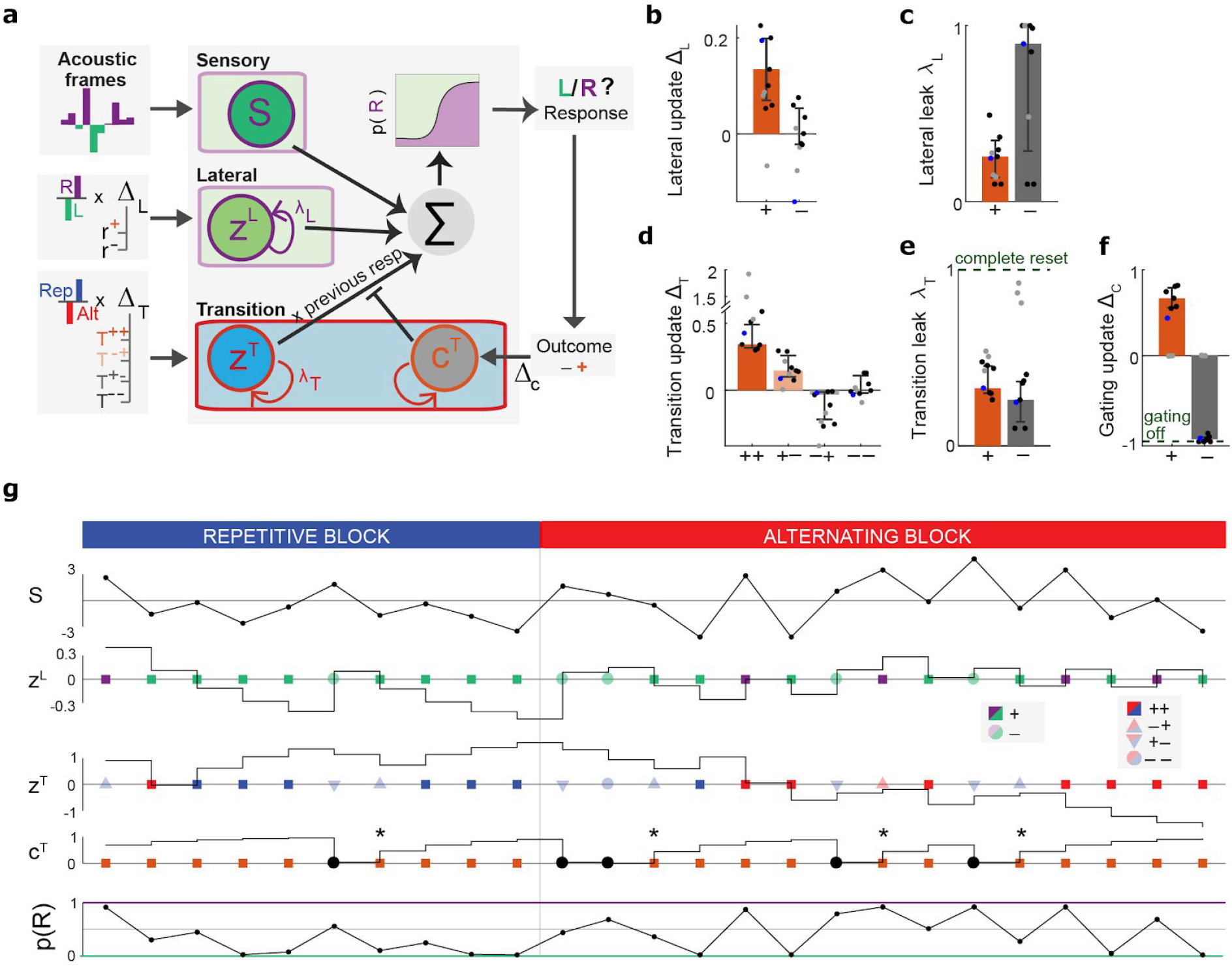
Dynamic generative model of history-dependent perceptual decisions. **a**, Architecture of the model. Decisions integrate the evidence from the sensory, lateral and transition modules. The sensory module accumulates the instantaneous evidence of each stimulus frame of the current trial. The lateral module maintains a representation of the lateral evidence *z*^*L*^ which is updated depending on the side and outcome of each trial response (updates ∆_L_), and on the outcome-dependent leak *λ*_L_. The transition module maintains the accumulated transition evidence *z*^*T*^, is updated depending on the last transition (i.e. ++,−+,+−,−−; updates ∆_T_) and on the outcome-dependent leak *λ*_T_. Evidence *z*^*T*^ is multiplied at each trial by the modulatory signal *c*^*T*^, updated based on the outcome of each response (updates ∆_C_), to yield the transition bias γ^T^= *z*^*T*^ × *c*^*T*^ × *r*_*t-1*_. The sum of sensory evidence and the lateral and transition biases determined the probability to choose either response at the current trial. Parameters were fitted to the choices of each rat separately. **b-e**, Best-fitting values of the update parameters in the generative model. Bars show median across seven rats (black and blue points). Three rats were excluded from the statistics because the fitted model yielded a solution without gating dynamics (gray points). **b**, lateral evidence update *∆*_*L*_ depending on trial outcome. **c**, outcome-dependent leak of the lateral bias *λ*_*L*_. **d**, transition evidence update *∆*_*T*_ depending on outcome of the last two trials (++,−+,+−,−−). **e**, outcome-dependent leak of the transition bias *λ*_*T*_. **f**, outcome-dependent update of the transition gating signal ∆_*C*_. A value of −1 correspond to an extinction of the gating signal on the subsequent trial (i.e. a full blockade of the corresponding bias), while +1 correspond to full recovery of the bias (i.e. gating equal to its maximum value of 1). **g**, Example of the model dynamics across 25 trials switching from a repetition to an alternation block (model parameters were fitted to an example animal, blue points in b-e). Traces from top to bottom depict the stimulus evidence *S*, z^*L*^, z^*T*^, *c*^*T*^ and overall probability to choose a Rightward response. The inputs to the variables *z*^*L*^, *z*^*T*^ and *c*^*T*^ are shown as a sequence of symbols on the corresponding trial axis: Left (green) vs. Right (purple) responses, repeating (blue) vs. alternating (red) transitions and rewarded (orange) vs. error (black) outcomes. Symbols shape represent different outcome combinations (see inset). Notice the reset of *c*^*T*^ after errors and the maintenance of *z*^*T*^ afterwards (asterisks).

We fitted the model parameters to the series of choices made by each rat (Fig. 6b-g and Supplementary Fig. 11) and obtained results in agreement with the gating hypothesis: first, correct transitions (++) led to strong changes in the transition evidence *z*^*T*^, while the other transitions (+−,−+,−−) did not lead to any consistent pattern (Fig. 6d). Second, the update parameters for *c*^*T*^ corresponded to a vanishing of this variable after errors for at least 7 rats out of 10, and a very strong recovery after any correct trial (Fig. 6f). This effectively converted the variable *c*^*T*^ into a gating variable that was able to completely block the use of the accumulated transition evidence *z*^*T*^ after a single error (Fig. 6g). By contrast, the leak of *z*^*T*^ was not significantly different after correct trials and after errors (p > 0.6, paired *t-*test, two-tailed), providing further evidence that the reset of the transition bias did not correspond to a loss of the accumulated evidence, as predicted by the Complete reset hypothesis (Fig. 5a top). Third, correct Rightward (Leftward) responses increased the lateral bias in favor of the Rightward (Leftward) response (Fig. 6c). Fourth, model comparison showed that this dynamical model gave a better account than versions where either *c*^*T*^ or the lateral bias *z*^*L*^ were omitted, as well as of the GLM described in the previous section (Supplementary Fig. 12). Finally, adding a modulatory variable *c*^*L*^ to the lateral module only had a marginal impact on model performance (Supplementary Fig. 13).

### Generative model simulation versus experimental data

Finally we assessed the capacity of the compact dynamical model to account for the dynamics of the previously reported repeating bias *b* (Fig. 2d-e) by comparing model simulations to actual rat data. The model very closely reproduced the build-up dynamics of *b* in series of correct repetitions and alternations (Fig. 7a). Moreover, the model allowed to partition the value of *b* into the contributions of the lateral and transition biases. While the transition bias was perfectly symmetric in series of repetitions and alternations (blue curves in Fig. 7a), the lateral bias behaved very differently: it only built up during series of repetitions, in which all the responses were on the same side, while it oscillated around zero in series of alternations, in which the contribution of each response was partially cancelled by the next one (green curves in Fig. 7a). Thus the dissection of the repeating bias into the lateral and transition biases explained the overall asymmetry found between the two blocks. Furthermore, the block asymmetry in the accumulation of the lateral bias also explained asymmetries in *b* found after correct unexpected responses (Supplementary Fig. 2b). Model simulations also reproduced the reset of repeating bias when a series of correct repetitions/alternations was interrupted by an error (Fig. 7b), and the subsequent rebound when the rat performed correctly again (Fig. 7c). Impressively, the model replicated the asymmetry in the magnitude of this rebound between the Repeating and Alternating blocks by summing (Fig. 7c top) or subtracting (Fig. 7c bottom), respectively, contributions of transition and lateral biases. Furthermore, the model provided a very good fit (Pearson’s r=0.96) to *b* for all possible sequences of 2-6 correct trials (Fig. 7d). In sum, by factorising the transition bias into accumulated transition evidence *z*^*T*^ and the modulatory signal *c*^*T*^, the model captured the non-linear across-trial dynamics of history-dependent biases pointing towards possible modulatory circuit mechanisms that could implement this computation (see Discussion).

**Figure 7.**
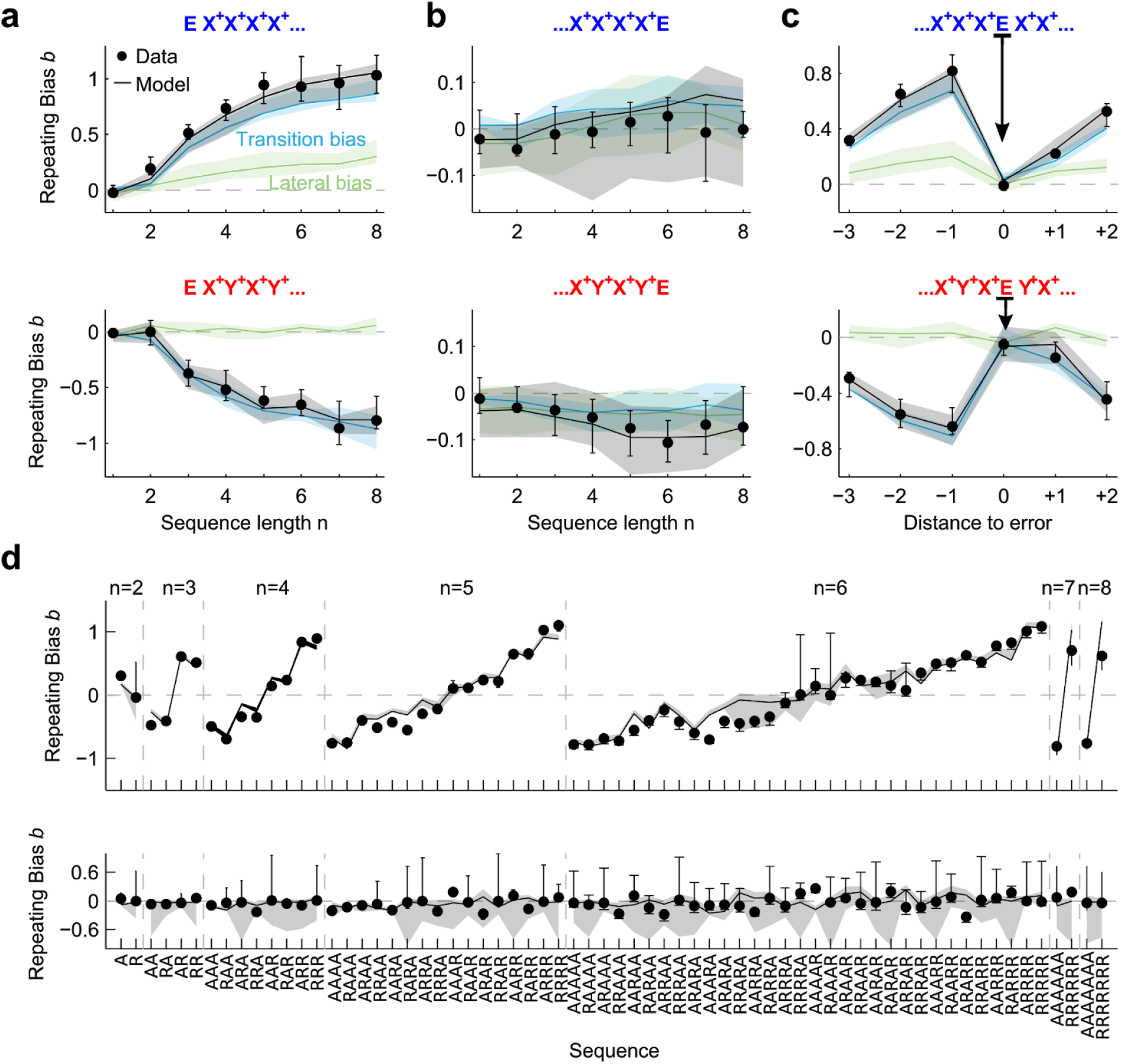
Generative model simulation compared to experimental data. Comparison between experimental data (dots) and model simulation (black curves) showing the Repeating bias *b* for different trial sequences. In the model, *b* was decomposed into the transition bias contribution (blue curves) and lateral bias contribution (green curves). **a**, Repeating bias *b* versus number *n* of correct repetitions (top) or alternations (bottom). Similar to color curves in Fig 2e. **b**, Repeating bias versus *n* after a repetitive (top) or alternating sequence (bottom) terminated by an error (as black curves in Fig. 2e). Notice the different range in the *b* axes compared with a. **c**, Repeating bias for sequences with an error E flanked by correct repetitions (top) or alternations (bottom). The bias *b* is given as a function the trial distance to the error response (distance zero represents *b* after the error). **d**, Repeating bias for all sequences made of *n* ≤ 8 repetitions (R) and alternations (A). Top panel shows correct sequences while bottom panel shows correct sequences terminated by an error. In all panels, data and model show median across *n*=10 rats. Error bars and shaded areas show first and third quartiles.

## DISCUSSION

We employed a standard acoustic 2AFC task to characterize how rats’ perceptual categorizations are affected by expectations derived from the history of past stimuli, choices and outcomes and how these expectations can be captured by a simple dynamical model. A thorough analysis of the behavior isolated two main sequential effects. First, we identified a sequential lateral effect that biased choices towards or away from the recently rewarded or unrewarded targets, respectively (Fig. 4a). This win-stay-lose-switch strategy has been extensively characterized both in humans (Abrahamyan et al., 2016; Braun et al., 2018; Fründ et al., 2014) and rodents (Akrami et al., 2018; Hwang et al., 2017). Second, we identified the sequential transition bias, a form of rule bias that had been previously shown to impact human reaction times (Cho et al., 2002; Kirby, 1976; Soetens et al., 1985), choices (Maloney et al., 2005) and neural responses (Jones et al., 2013; Sommer et al., 1999). Our results however go beyond from previous reports in several important aspects regarding error responses: first, repetitions or alternations did not influence subsequent choices whenever one of the two trials of the transition was unrewarded, meaning that the running estimate of the transition probabilities only accumulated evidence from repetitions and alternations of two rewarded trials. Second, the transition bias was reset after an error trial, i.e. animal responses temporarily ignored the recent history of rewarded repetitions and alternations. However, this reset did not imply the reset of the accumulated transition evidence, i.e. the tallie keeping track of the number of recent repetition vs. alternations, whose influence over behavior was restored as soon as the animal obtained a reward (Fig. 5c).

The probability of a subject to repeat the previous response (Fig. 7a) is a common measure to characterize history effects (Abrahamyan et al., 2016; Urai et al., 2018). By dissecting the distinct contribution of both first- and second-order serial biases (Gokaydin et al., 2011; Jones et al., 2013; Meyniel et al., 2016; Wilder et al., 2009), i.e. the lateral and the transition biases, respectively, to the repeating bias we were able to understand the asymmetry in its magnitude between the Repeating and Alternating blocks (Fig. 2c-e): in series of correct repetitions, both transition and lateral bias add up and yield a strong tendency of the animals to repeat the last rewarded response (Fig. 7a top). In contrast, in alternating environments, the lateral bias does not build up and the negative repeating bias (tendency to switch) is solely given by the transition bias (Fig. 7a bottom). In sum, the first-order lateral bias favors repetition over alternation; the second-order transition bias has a symmetric effect. In fact, our analysis provides indirect evidence that animals recapitulated previous repetitions and alternations into a single and symmetric transition bias and not into separate variables (Supplementary Fig. 7c). A recent modeling study has proposed that estimating first and second-order rates is part of the same core computation that the brain performs when analyzing binary sequences. This computation comes down to estimate the two independent transition probabilities P(L_*t*_|R_*t*-1_) and P(R_*t*_|L_*t*-1_) between consecutive trials *t* −1 and *t* (Meyniel et al., 2016). Our findings seem at odds with this hypothesis because the dependence of each type of bias on the response outcome was very different: whereas incorrect responses *r*^*−*^ tended to cause a negative switch effect (Fig. 4a), incorrect transitions (*T*^+−^,*T*^*−+*^ and *T*^*−−*^) had no effect (Fig. 4b). Furthermore, only the transition bias showed a reset-and-rebound dynamics caused by error responses (Fig. 5c and Supplementary Fig. 5g,h). An alternative hypothesis, based on analysis of response evoked potentials (ERP), proposes that the lateral bias is generated by the *processing of the response* whereas the transition bias from the *stimulus processin*g (Jones et al., 2013; Wilder et al., 2009). Preliminary data obtained in the same task in the absence of any stimuli seems to indicate that the transition bias is still present and thus does not seem to be contingent on the processing of sensory inputs.

Several of our findings, together with previous literature (Cho et al., 2002; Jones et al., 2013; Kirby, 1976; Soetens et al., 1985; Sommer et al., 1999), suggest that the transition bias is a fundamental aspect of sequence processing preserved across subjects, species and conditions and which does not seem particularly adaptive to the details of the experiment. First, the transition bias was the same in both Repeating and Alternating blocks (Supplementary Fig. 8) reflecting the use of a single fixed strategy that could switch from generating a net positive repeating bias in a Repeating block to generating a negative bias in the alternating block (Fig. 2e). Interestingly, this invariance of the transition bias across the Repetitive and Alternating blocks has also been found in humans performing a 2AFC task (Jones et al., 2013). Second, the transition bias was also present when sequences are uncorrelated and the bias can only hinder performance (Supplementary Fig. 10c; (Cho et al., 2002; Jones et al., 2013; Kirby, 1976; Soetens et al., 1985; Sommer et al., 1999). Third, the trial integration window over which animals estimated the repetition rate (~3-5 trials; Fig. 4b) does not seem adapted to the block length (200 trials). This short-span estimate allowed to reverse the repetition bias rapidly after a block switch (Supplementary Fig. 1a) at the cost of a noisier estimate of the repetition rate (Gallistel et al., 2004; Nassar et al., 2010; Sutton and Barto, 2018). Quantification of this integration window in human subjects performing different 2AFC tasks yields numbers in the range of 2-10 trials, despite the use of very long trial-blocks with constant sequence correlations (Jones et al 2013). Thus, rather than an overestimation of the environment’s volatility (Behrens et al., 2007; Nassar et al., 2010), the short fixed windows might reflect structural adaptation to the statistics of natural environments (Seriès and Seitz, 2013) or a capacity limitation of the system. Fourth, the sophisticated outcome-dependent across-trial dynamics of the transition bias were found systematically in every animal we tested (Fig. 4c) showing that they do not reflect idiosyncratic strategies but the action of an unknown basic cognitive process. Finally, there was one aspect of the mechanism that seemed adaptive: the magnitude of the transition kernel gradually increased when animals, initially trained using uncorrelated sequences, were presented with correlated sequences (Supplementary Fig. 10). Rats in fact had been previously shown to suppress sequential biases when those can be turned up against them (Tervo et al., 2014). Thus, the transition bias can be adapted to the temporal structure of the environment, if not in nature, at least in magnitude (Abrahamyan et al., 2016) (Supplementary Fig. 10).

Why does the transition bias reset after errors? Previous studies have shown that an uncued change in stimulus-outcome contingencies leading to an unexpected number of unrewarded choices can trigger an abrupt behavioral change in rats, switching from the exploitation of a statistical model of the environment to an exploration mode in which they sample the environment in an unbiased way in order to build new beliefs (Karlsson et al., 2012). This suggests that the reset-and-rebound dynamics of the transition bias could be interpreted as a fast switching between the exploitation of their internal model, represented by their estimate of the transition probability, to a mode that relies almost exclusively on sensory information. This expectation-free mode, however, is different from the standard exploration mode in which animals guide their choices aiming to reduce the uncertainty of the environment. In contrast, our animals, perhaps unable to use their prior after not obtaining the reward (i.e. not knowing *what* they must repeat/alternate after an error), guide their choices based on the sensory evidence alone. To capture the reset-and-rebound dynamics, we built a generative-sufficient novel model that could jointly describe the latent trial-to-trial dynamics of (1) the expectation formation following standard reinforcement learning updating rules (Behrens et al., 2007) (Fig. 6a-e) and (2) a modulatory signal *c*^*T*^ that had a multiplicative effect on the impact of the transition evidence in biasing choices. The fitting of the model parameters revealed that *c*^*T*^ reset to zero after errors and then increased progressively with a series of correct trials (Fig. 6f). This modulatory variable may reflect subjects’ confidence in their internal model of the environment statistics or, alternatively, the probability that the subject operated in the exploitation mode versus the expectation-free mode. Furthermore, in this expectation-free mode in which the prior is not used, it also cannot be updated with new transition information, as can be concluded from the finding that only ++ transitions impacted subsequent choices (Fig. 4b).

Recent studies found that, in the absence of feedback, the impact of a choice on the subsequent trial was weaker if the subject was unsure of her choice (Braun et al., 2018; Samaha et al., 2018; Urai et al., 2017). The explanation provided in two of these studies was that, according to a normative theory describing how to accumulate noisy evidence in the face of uncued changes (Glaze et al., 2015), low confidence choices should have a weaker contribution on the belief about what will be the next stimulus category (Braun et al., 2018). In our latent variable model, this is is indeed true because unrewarded transitions, *T*^*+−*^ and *T*^*−−*^, supposedly generating the lowest confidence about what the true transition was, have a weaker contribution to the accumulated evidence *z*^*T*^ (see fitted values of *Δ*_*T*_ in Fig. 6d). However, a bias reset after incorrect or low confidence trials was not reported in these studies, i.e. errors without feedback did not seem to modulate retroactively the impact of previous trials onto the next choice, unlike what was observed in our rats. Also, in (Braun et al., 2018) subjects were informed about the existence of “more repeating”, “more alternating” and “uncorrelated” sessions. In contrast, our animals were constantly estimating the transition probability which varied in blocks during each session. A fair assessment on the existence of the expectation bias reset in humans would necessitate of an experiment in which subjects are blind about the sequence correlations but receive feedback in every trial.

The activation of noradrenergic inputs onto the anterior cingulate cortex has been shown to control the switching into a behavioral mode in which beliefs based on previous experience do not guide choices (Tervo et al., 2014). Because in the quoted study the experimental condition was a free choice task, removing the impact of history-effects resulted in stochastic exploration (Tervo et al., 2014). This prompts the question of whether the activation of the very same modulatory pathway underlies the after-error switch into the expectation-free sensory-based mode observed in our task. Future pharmacological an electrophysiological experiments will shed light into the brain regions encoding the expectation signals, their modulatory variables as well the circuit mechanisms underlying their combination with the incoming sensory information.

## METHODS

All experimental procedures were approved by the local ethics committee (Comité d’Experimentació Animal, Universitat de Barcelona, Spain, Ref 390/14).

### Animal Subjects

Animals were male Long-Evans rats (n=25, 350-650g; Charles River), pair-housed during behavioral training and kept on stable conditions of temperature (23°C) and humidity (60%) with a constant light-dark cycle (12h:12h, experiments were conducted during the light phase). Rats had free access to food, but water was restricted to behavioral sessions. Free water during a limited period was provided on days with no experimental sessions.

### Task description

The two tasks performed were auditory reaction-time two-alternative forced choice procedures: an LED on the center port indicated that the rat could start the trial by poking in (Fig. 1a). After a fixation period of 300 ms, an acoustic stimulus consisting in a superposition of two amplitude-modulated sounds (see details below) was presented. The rats had to discriminate the dominant sound and seek reward in the associated port. Animals could respond any time after stimulus onset. Withdrawal from the center port during the stimulus immediately stopped the stimulus. Correct responses were rewarded with a 24 µl drop of water and incorrect responses were punished with a bright light and a 5 s time-out. Trials in which the rat did not make a side poke response within 4 seconds after leaving the center port were considered invalid trials and were excluded from the analysis (on average, only 0.4% of the trials were invalid). Behavioral setup (Island Motion, NY) was controlled by a custom software developed in Matlab (Mathworks, Natick, MA), based on the open-source BControl framework (http://brodylab.princeton.edu/bcontrol). Rats performed an average of 694 trials per session (range: 335 - 1188), one session per day lasting 60-90 min, 6 days per week, during 9 months. Rats were trained using an automated training protocol that had several stages and lasted between 2 and 3 months (depending of the animal). The data presented in this study was taken from the period after training yielding an average of 56,506 valid trials per rat. A first group of ten rats were trained in the frequency discrimination version of the task (see below) in which the correlated sequence of trials was present from the training. A subset of three rats from this group were also trained in a random time-out version of the task where the duration of the after-error time-out was randomly chosen between 1, 3 or 5 s. A second group of n=9 rats were trained in the same frequency discrimination version of the task but starting with uncorrelated stimulus sequences and only after several weeks, introducing the correlated sequences used in the first group of animals. A third group of n=6 rats were trained in a level discrimination version of the task using the same correlated sequence than the first group.

### Acoustic stimulus

In the two acoustic tasks used, the stimulus *S*(*t*) in each trial was created by simultaneously playing two amplitude modulated (AM) sounds *s*_*R*_(t) and *s*_*L*_(t):

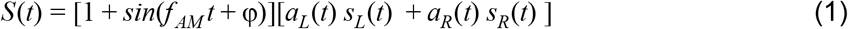

The AM frequency was *f*_AM_=20 Hz and the phase delay φ = 3π/2 made the envelope zero at *t*=0. In the frequency discrimination task, *s*_*L*_(t) and *s*_*R*_(t) were pure tones with frequencies 6.5 kHz and 31 kHz, respectively, played simultaneously in the two speakers. In the level discrimination task (Supplementary Fig. 9) they were broadband noise played either from the Left or the Right speaker, respectively. The amplitudes of the sounds *s*_L_(*t*) and *s*_R_(*t*) were separately calibrated at 70 dB. Sounds were delivered through generic electromagnetic dynamic speakers (STAX, SRS-2170) located on each side of the chamber, and calibrated using a free-field microphone (Med Associates Inc, ANL-940-1).

### Stimulus Sequence

The Markov chain generated the sequence of stimulus category *c*_*k*_= {-1,1}, that determined whether the reward in the *k*-th trial was available in the Left or the Right port respectively (Fig. 1b top). The stimulus category set which of the two sounds *s*_*L*_(t) and *s*_*R*_(t) composing each stimulus was dominant, which ultimately determined the statistics of the sound amplitudes *a*_1_(t) and *a*_2_(t) (Eq. 1) as described below. In each trial, independently of *c*_*k*_, the stimulus strength *s*_*k*_ was also randomly generated (Fig. 1b bottom). Stimulus strength defined the relative weights of the dominant and non-dominant sounds: for example, when *s*_*k*_ =1 only the dominant sound was played (i.e. easiest trials) whereas when *s*_*k*_ =0 the two sounds had on average the same amplitude (i.e. hardest trials). We used four possible values for *s* = 0, 0.23, 0.48 and 1. The stimulus evidence was defined in each trial as the combination *e*_*k*_ = *c*_*k*_\**s*_*k*_. The value of *e*_*k*_ determined the p.d.f. from which the instantaneous evidence *x*(*t*) was drawn in each frame *t* (i.e. in each 50 ms AM-envelope cycle; Fig. 1d top): when *e*_*k*_:=±1 the p.d.f. was *f* (*x*) = δ(*x*∓1) (i.e. a Dirac delta p.d.f.) whereas when *e*_*k*_ ∈ (−1,1), it was a stretched Beta distribution with support [-1,1], mean equal to *e*_*k*_ and variance equal to 0.06 (Fig. 1d top). Finally, the amplitudes *a*_*L*_(*t*) and *a*_*R*_(*t*) of the two AM envelopes (Eq. 1) were obtained using *a*_*L*_(*t*)=[1+*x*(t)]/2 and *a*_*R*_(*t*)=[1-*x*(*t*)])/2 (see example in Fig. 1d). With this choice the sum of the two envelopes was constant in all frames *a*_*L*_(*t*)+*a*_*R*_(*t*)=1.

### Psychometric curve analysis

We computed two types of psychometric curves for each animal, by pooling together trials across all sessions for each type of block and for each of the 7 different stimulus evidences (*e*= 0, ±0.23, ±0.48, ±1). We calculated (1) the proportion of Rightward responses vs. stimulus evidence *e* (Fig. 1a left) and (2) the Proportion of Repeated responses as function of Repeating Stimulus Evidence (Fig. 1b), where positive or negative Repeating Stimulus Evidence denote trials in which the animals had evidence to repeat their previous choice (e.g. a Rightward stimulus with evidence *e =* +0.23 after a Left response implied a repeating stimulus evidence equal to - 0.23). Both psychometric curves were separately fitted to a 2-parameter probit function (using Matlab function *nlinfit*):

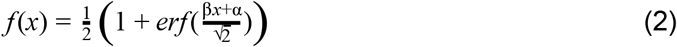

The sensitivity *β* quantified the stimulus discrimination ability, while the bias *α* in the Proportion of Rightward responses captures the animal fixed side preference for the Left or Right port, and the bias *α* in the Proportion of Repeated responses arose from the expectation generated by the history of recent choices, i.e. it showed the animal’s tendency to repeat or alternate their previous choice. We termed these two biases the fixed side bias *B* and the repeating bias *b*, respectively. Within-subject error bars were estimated by one standard deviation of a non-parametric bootstrap (n=1000). Across-subject error bars, corresponded to the 1^st^ and 3^rd^ quartiles.

## ACKNOWLEDGEMENTS

We thank Anke Braun, Anne E. Urai, Daniel Duque, Gabriela Mochol, Genís Prat-Ortega, Lluís Hernández, Manuel Molano, Tobias H. Donner and Yerko Fuentealba for excellent discussions and critical reading of the manuscript and Lejla Bektic for help with training of the animals. This research was supported by the Spanish Ministry of Economy and Competitiveness together with the European Regional Development Fund (BES-2011-049131 to A.H.M.; SAF2013-46717-R, SAF2010-15730, RYC-2009-04829 and SAF2015-70324-R to J.R.), the European Research Council (ERC-2015-CoG - 683209 Priors to J.R; ERC-2013-CoG - 615699 Neurokinematics to D.R), the European Union (Marie Curie IRG PIRG07-GA-2010-268382 to J.R.; Marie Curie auction CEMNET project 629613 to A.H.; Marie Curie IIF253873 to P.R.-O.), Mexico (CONACyT-144335 to P.R.-O.) and USA (National Institute on Deafness and Other Communication Disorders-R01DC015531 to S.J.).

## AUTHOR CONTRIBUTIONS

A.H.M. and J.R. designed the experiments with the assistance from D.R. P.R.-O. and S.J.; A.H.M. carried out all the experiments; A.H.M. and A.H. analyzed the data; A.H. developed the generative dynamical model; A.H.M., A.H. and J.R. interpreted the data and wrote the manuscript with contributions from the rest of the authors.

## REFERENCES

Abrahamyan, A., Silva, L.L., Dakin, S.C., Carandini, M., and Gardner, J.L. (2016). Adaptable history biases in human perceptual decisions. Proc. Natl. Acad. Sci. U. S. A. 113, E3548–E3557.

Akaishi, R., Umeda, K., Nagase, A., and Sakai, K. (2014). Autonomous mechanism of internal choice estimate underlies decision inertia. Neuron 81, 195–206.

Akrami, A., Kopec, C.D., Diamond, M.E., and Brody, C.D. (2018). Posterior parietal cortex represents sensory history and mediates its effects on behaviour. Nature 554, 368–372.

Ashourian, P., and Loewenstein, Y. (2011). Bayesian inference underlies the contraction bias in delayed comparison tasks. PLoS One 6, e19551.

Behrens, T.E.J., Woolrich, M.W., Walton, M.E., and Rushworth, M.F.S. (2007). Learning the value of information in an uncertain world. Nat. Neurosci. 10, 1214–1221.

Braun, A., Urai, A.E., and Donner, T.H. (2018). Adaptive History Biases Result from Confidence-weighted Accumulation of Past Choices. J. Neurosci.

Busse, L., Ayaz, A., Dhruv, N.T., Katzner, S., Saleem, A.B., Schölvinck, M.L., Zaharia, A.D., and Carandini, M. (2011). The detection of visual contrast in the behaving mouse. J. Neurosci. 31, 11351–11361.

Cho, R.Y., Nystrom, L.E., Brown, E.T., Jones, A.D., Braver, T.S., Holmes, P.J., and Cohen, J.D. (2002). Mechanisms underlying dependencies of performance on stimulus history in a two-alternative forced-choice task. Cogn. Affect. Behav. Neurosci. 2, 283–299.

Corrado, G.S., Sugrue, L.P., Seung, H.S., and Newsome, W.T. (2005). Linear-Nonlinear-Poisson models of primate choice dynamics. J. Exp. Anal. Behav. 84, 581–617.

Daw, N.D., O’Doherty, J.P., Dayan, P., Seymour, B., and Dolan, R.J. (2006). Cortical substrates for exploratory decisions in humans. Nature 441, 876–879.

Ebitz, R.B., Albarran, E., and Moore, T. (2018). Exploration Disrupts Choice-Predictive Signals and Alters Dynamics in Prefrontal Cortex. Neuron 97, 475.

Ernst, M.O., and Banks, M.S. (2002). Humans integrate visual and haptic information in a statistically optimal fashion. Nature 415, 429–433.

Fischer, J., and Whitney, D. (2014). Serial dependence in visual perception. Nat. Neurosci. 17, 738–743.

Fründ, I., Wichmann, F.A., and Macke, J.H. (2014). Quantifying the effect of intertrial dependence on perceptual decisions. J. Vis. 14.

Gallistel, C.R., Fairhurst, S., and Balsam, P. (2004). The learning curve: implications of a quantitative analysis. Proc. Natl. Acad. Sci. U. S. A. 101, 13124–13131.

Glaze, C.M., Kable, J.W., and Gold, J.I. (2015). Normative evidence accumulation in unpredictable environments. Elife 4.

Gokaydin, D., Ma-Wyatt, A., Navarro, D., and Perfors, A. (2011). Humans use different statistics for sequence analysis depending on the task. (Cognitive Science Society),.

Goldfarb, S., Wong-Lin, K., Schwemmer, M., Leonard, N.E., and Holmes, P. (2012). Can post-error dynamics explain sequential reaction time patterns? Front. Psychol. 3, 213.

Hwang, E.J., Dahlen, J.E., Mukundan, M., and Komiyama, T. (2017). History-based action selection bias in posterior parietal cortex. Nat. Commun. 8, 1242.

Jones, M., Curran, T., Mozer, M.C., and Wilder, M.H. (2013). Sequential effects in response time reveal learning mechanisms and event representations. Psychol. Rev. 120, 628–666.

Karlsson, M.P., Tervo, D.G.R., and Karpova, A.Y. (2012). Network resets in medial prefrontal cortex mark the onset of behavioral uncertainty. Science 338, 135–139.

Kim, T.D., Kabir, M., and Gold, J.I. (2017). Coupled Decision Processes Update and Maintain Saccadic Priors in a Dynamic Environment. J. Neurosci. 37, 3632–3645.

Kirby, N.H. (1976). Sequential effects in two-choice reaction time: automatic facilitation or subjective expectancy? J. Exp. Psychol. Hum. Percept. Perform. 2, 567–577.

Lau, B., and Glimcher, P.W. (2005). Dynamic response-by-response models of matching behavior in rhesus monkeys. J. Exp. Anal. Behav. 84, 555–579.

Maloney, L.T., Dal Martello, M.F., Sahm, C., and Spillmann, L. (2005). Past trials influence perception of ambiguous motion quartets through pattern completion. Proc. Natl. Acad. Sci. U. S. A. 102, 3164–3169.

Meyniel, F., Maheu, M., and Dehaene, S. (2016). Human Inferences about Sequences: A Minimal Transition Probability Model. PLoS Comput. Biol. 12, e1005260.

Nassar, M.R., Wilson, R.C., Heasly, B., and Gold, J.I. (2010). An approximately Bayesian delta-rule model explains the dynamics of belief updating in a changing environment. J. Neurosci. 30, 12366–12378.

Nogueira, R., Abolafia, J.M., Drugowitsch, J., Balaguer-Ballester, E., Sanchez-Vives, M.V., and Moreno-Bote, R. (2017). Lateral orbitofrontal cortex anticipates choices and integrates prior with current information. Nat. Commun. 8, 14823.

Pardo-Vazquez, J.L., Castineiras, J., Valente, M., and Costa, T. (2018). Weber’s law is the result of exact temporal accumulation of evidence. bioRxiv.

Samaha, J., Switzky, M., and Postle, B.R. (2018). Confidence boosts serial dependence in orientation estimation. bioRxiv.

Seriès, P., and Seitz, A.R. (2013). Learning what to expect (in visual perception). Front. Hum. Neurosci. 7, 668.

Soetens, E., Boer, L.C., and Hueting, J.E. (1985). Expectancy or automatic facilitation? Separating sequential effects in two-choice reaction time. J. Exp. Psychol.

Sommer, W., Leuthold, H., and Soetens, E. (1999). Covert signs of expectancy in serial reaction time tasks revealed by event-related potentials. Percept. Psychophys. 61, 342–353.

Stocker, A.A., and Simoncelli, E.P. (2006). Noise characteristics and prior expectations in human visual speed perception. Nat. Neurosci. 9, 578–585.

Sutton, R.S., and Barto, A.G. (2018). Reinforcement Learning: An Introduction (MIT Press).

Tervo, D.G.R., Proskurin, M., Manakov, M., Kabra, M., Vollmer, A., Branson, K., and Karpova, A.Y. (2014). Behavioral variability through stochastic choice and its gating by anterior cingulate cortex. Cell 159, 21–32.

Urai, A.E., Braun, A., and Donner, T.H. (2017). Pupil-linked arousal is driven by decision uncertainty and alters serial choice bias. Nat. Commun. 8, 14637.

Urai, A.E., de Gee, J.W., Tsetsos, K., and Donner, T.H. (2018). Choice history biases subsequent evidence accumulation.

Vernon, G., Farrow, D., and Reid, M. (2018). Returning Serve in Tennis: A Qualitative Examination of the Interaction of Anticipatory Information Sources Used by Professional Tennis Players. Front. Psychol. 9, 895.

Wilder, M., Jones, M., and Mozer, M.C. (2009). Sequential effects reflect parallel learning of multiple environmental regularities. In Advances in Neural Information Processing Systems 22, Y. Bengio, D. Schuurmans, J.D. Lafferty, C.K.I. Williams, and A. Culotta, eds. (Curran Associates, Inc.), pp. 2053–2061.

Yu, A.J., and Cohen, J.D. (2008). Sequential effects: Superstition or rational behavior? Adv. Neural Inf. Process. Syst. 21, 1873–1880.

