## Supplemental Material for "Response outcomes gate the impact of expectations on perceptual decisions"

Supplementary Materials:

Supplementary Figures ........................................................pag. 1

Supplementary Methods .................................................... pag. 13

#### Supplementary Figures

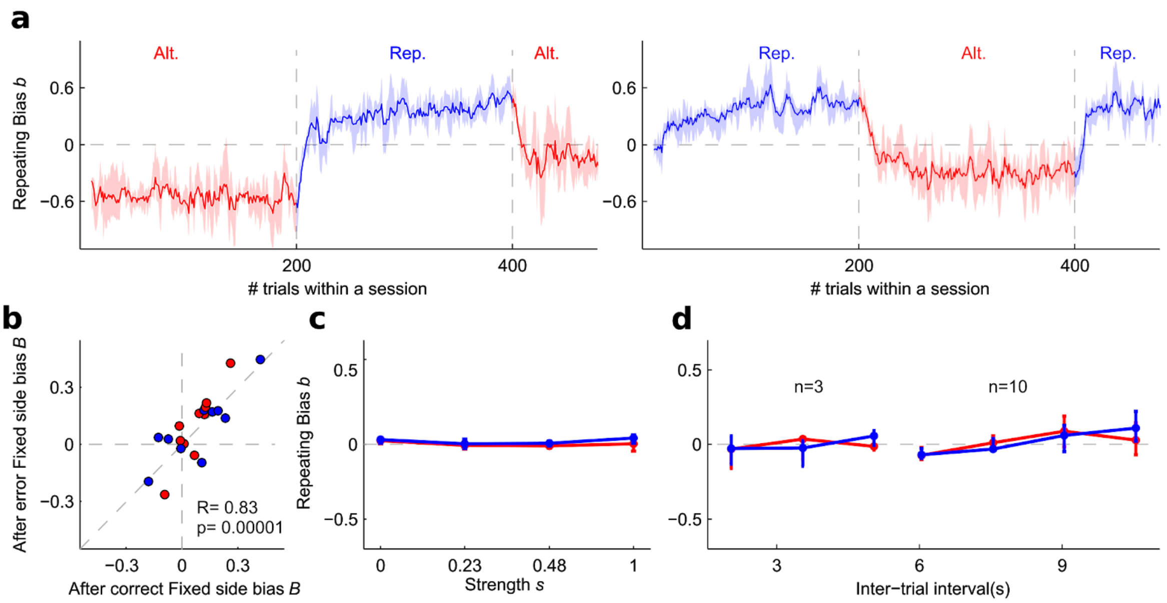


**Supplementary Figure 1. Time course of Repeating Bias across trials. a,** Time course of repeating bias *b* after correct responses through the course of one session starting with an Alternating block (left) or a Repetitive block (right). Repetitive blocks and Alternating blocks are indicated in blue and red color, respectively (vertical lines indicate block transitions). Curves show median *b* over *n* = 9 rats (one rat was excluded from the analysis because it only completed about 258 trials per session) computed using a 10 trials sliding window. The shaded areas illustrate the 1st and 3rd quartiles. **b,** Fixed side bias *B* after error trials versus *B* after correct trials. Each dot represent one animal (*n* = 10) in Repetitive block (blue) or Alternating Block (red). **c,** Repeating bias *b* after error trials versus previous stimulus strength in the Alternating (red) and Repetitive blocks (blue). Dots show median across *n* = 10 animals and error bars the 1st and 3rd quartiles. **d,** Repeating Bias *b* after error trials versus inter-trial interval (ITI) in the Alternating (red) and Repetitive blocks (blue) for rats punished with a fixed 5 s after-error time-out (*n* = 10, right) or with a randomly interleaved 1, 3 or 5 s time-out (*n* = 3, left). Dots show median across *n* = 3 (right) and *n* = 10 (left) animals and error bars the 1st and 3rd quartiles.

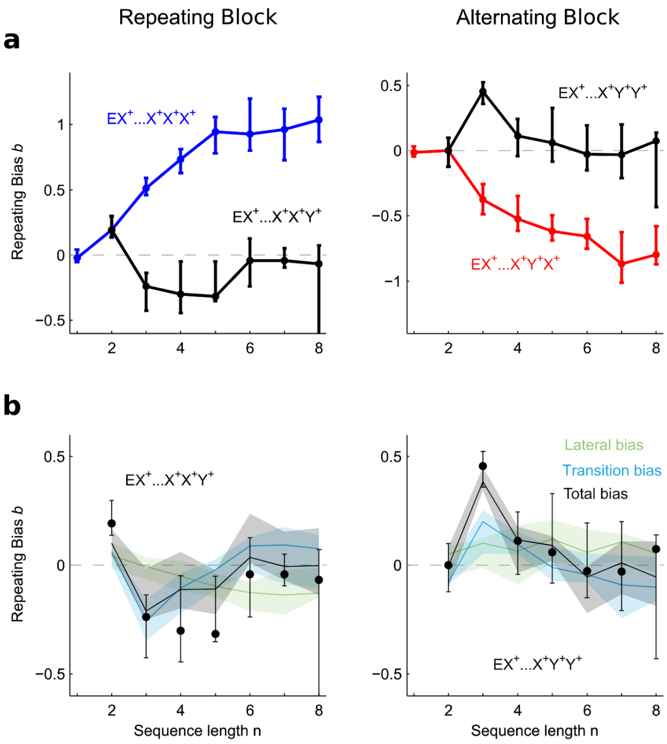


**Supplementary Figure 2. Repeating bias after unexpected correct responses. a,** Repeating bias *b* versus the number of correct previous repetitions in Repetitive block (blue) or alternations in Alternating block (red). Black dots show *b* after one unexpected correct alternation following several repetitions (left) or an unexpected correct repetition following several alternations (right). Breaking the blocks common pattern sequence with correct unexpected response didn't reset the repeating bias (compare black dots with those in Fig. 2e). **b,** Comparison between experimental data (dots) and latent variable model simulation (black curves) for the repeating bias *b* after one unexpected correct responses (same as black dots in **a**). In the model, *b* was decomposed into the transition bias (blue curves) and lateral bias (green curves). Dots show median across *n* = 10 animals (Group 1). Error bars and shaded areas show the 1st and 3rd quartiles.

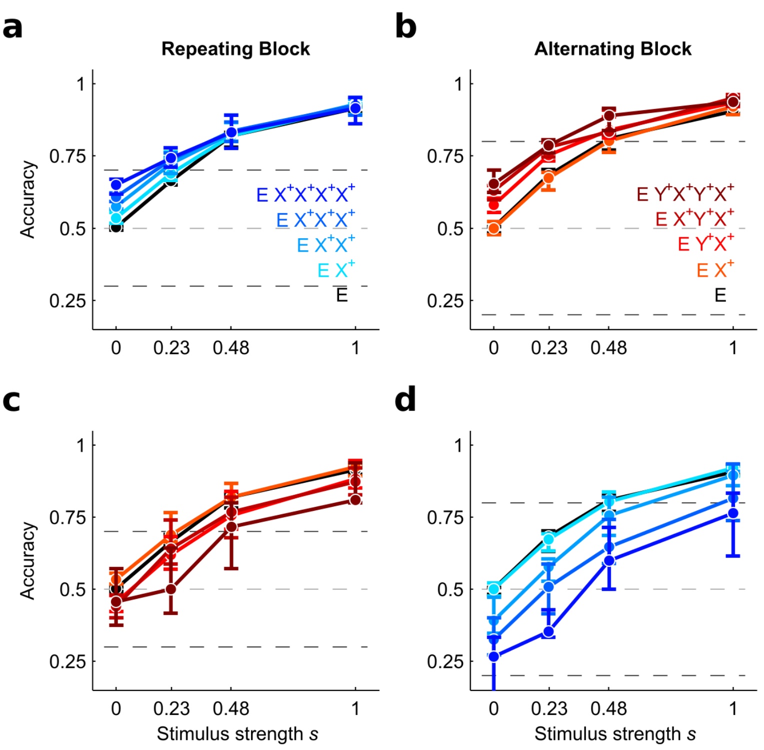


**Supplementary Figure 3. The repeating bias build-up from previous trials impacts the rats' classification accuracy.** Accuracy, defined as the fraction of correct responses, versus the current stimulus strength $s_k$ in trials following several correct repetitions (**a,d**; blue gradient, see inset for color codes) or alternations (**b,c**; red gradient). We separately computed the accuracy after each sequence for the Repetitive block (**a, c**) or for the Alternating block (**b, d**). Accuracy increased when the past sequence yielded a repeating bias that was congruent with the block's tendency (**a-b**) and decreased when it was incongruent (**c-d**). The impact of the repeating bias on accuracy was most noticeable on stimuli with low stimulus strength. Dashed lines mark the maximum and minimum performance that can be reached for stimuli with $s$ = 0 in each block if subjects adopted the strategy of always repeating or alternating the previous response. In the repeating block these values are the sequence repeating probability $P_{\text{rep}}$ and 1- $P_{\text{rep}}$, respectively, whereas in the alternating block they are 1- $P_{\text{rep}}$ and $P_{\text{rep}}$. Dots show median across $n$ = 10 animals (Group 1). Error bars and shaded areas show the 1st and 3rd quartiles.

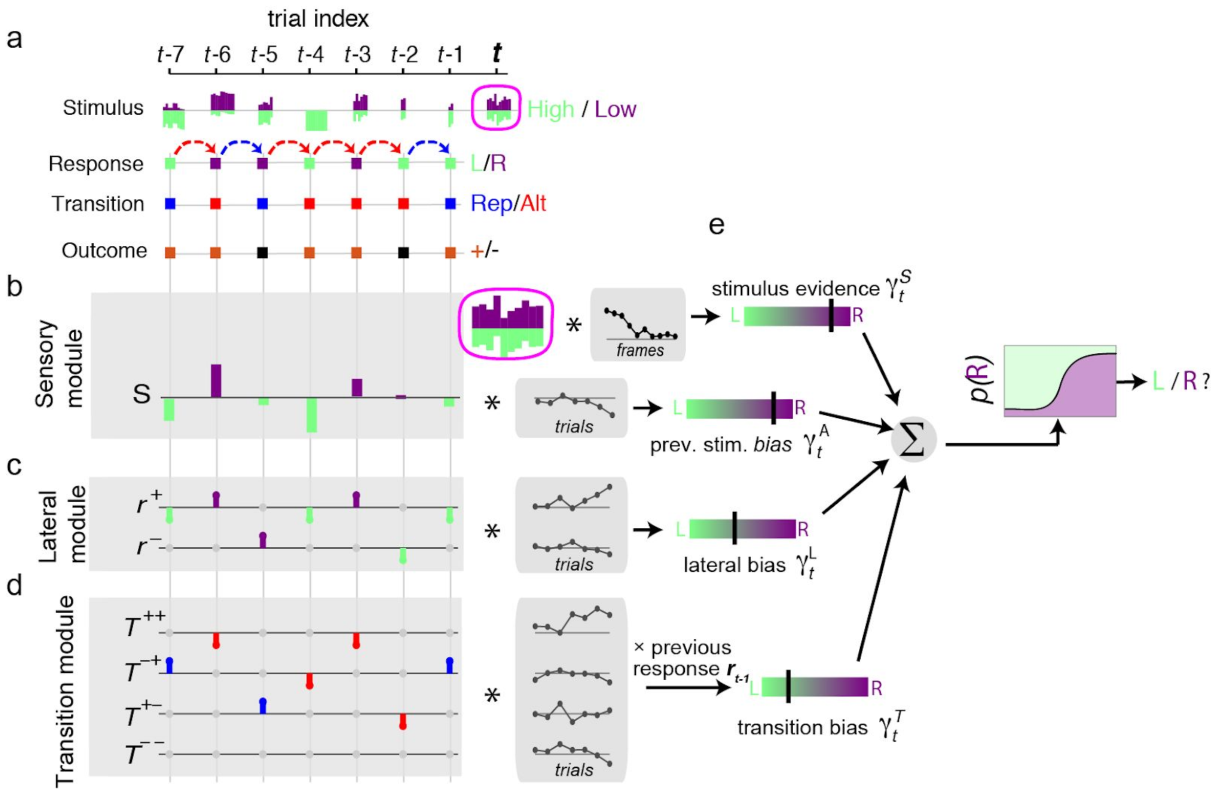


**Supplementary Figure 4. GLM assessing the impact of sensory evidence and recent history onto rats choices**. **a,** Exemplar series of recent history trials used to model rat decisions at current trial *t*. The Stimulus series depicts the amplitude of each acoustic frame for the low (green bars) and high frequency tones (purple bars). The Response series shows the rat decisions (green squares for Leftward and purple squares for Rightward choices). The Transition series shows the relation of two consecutive responses (blue for Repetition, red for Alternation) and the Outcome series indicates whether the responses were rewarded (orange) o not (black). These series are combined to generate the regressors that are grouped into the Sensory (b), Lateral (c) and Transition (d) modules. **b,** The amplitude difference of each acoustic frame of the current stimulus (pink box) is weighted separately by a stimulus kernel which is fitted for each animal (black dots). The outcome of this sum provides the Stimulus evidence $\gamma^S{}_t$. which can take values ranging from strong Left to strong Right evidence (color bar with green-purple gradient; black tick shows the value of the example sequence). The net stimulus evidence of each of the previous trials (*t-1*, *t-2*, …, green and purple bars in gray box) are weighted by the previous stimulus kernel providing the After-effect bias $\gamma^A{}_t$. **c,** The Lateral module weights separately previous rewarded $r^+$ and unrewarded responses $r^-$ that take the values -1 (Left), +1 (Right) or 0. The sum of these two series gives rise to the lateral bias $\gamma_t^L$. **d,** Transitions are considered separately depending on the outcome of the two trials in the transition: $T^{++}$ (rewarded-rewarded), $T^{-+}$ (error-rewarded), $T^{+-}$ (rewarded-error) and $T^{--}$ (error-error) which take the values -1 (alternation), +1 (repetition) and 0. The weighted sum of transition regressors is then multiplied by the previous response $r_{t-1}$ in order to yield the transition bias $\gamma^T$ which provides Rightward vs Leftward evidence. **e,** The sum $\gamma^S{}_t + \gamma^A{}_t + \gamma^L{}_t + \gamma^T{}_t$ is then passed through a sigmoid function to yield the trial-by-trial probability of selecting a Right response. For each rat, the weights of the kernels (shown as black connected dots in the center gray boxes) were fitted to maximize the model's probability to generate the actual animal choices.

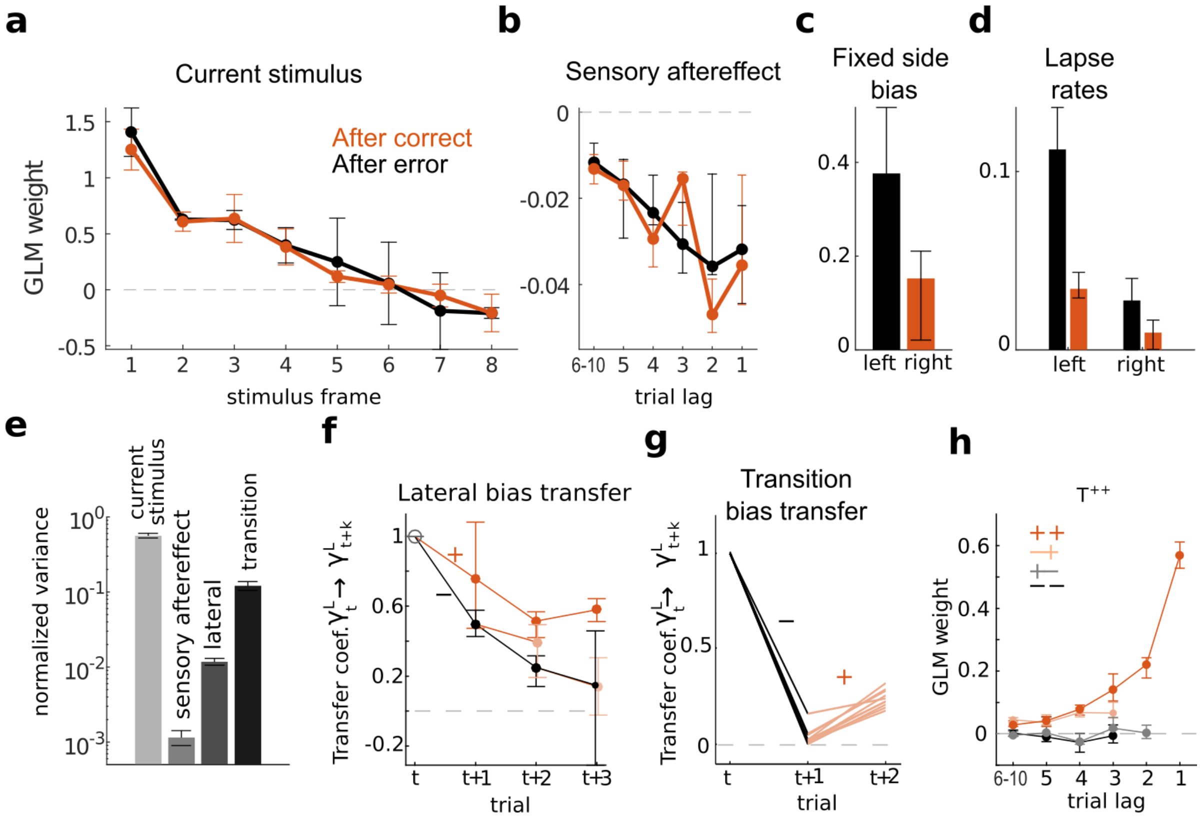


**Supplementary Figure 5. Fitted weights quantifying the impact onto animals decisions of the different modules.** Average coefficients ($n$ = 10 animals) obtained in the GLM when separately fitting the choices in trials after a correct (orange) and error response (black). **a,** Frame-by-frame influence of the current stimulus onto rats decisions, commonly termed psychophysical kernel, shows that the current stimulus, and in particular the first acoustic frames in the stimulus, had a strong impact on decisions. **b,** Influence of the sensory after-effect caused by the net sensory evidence of previous trials as a function of trial lag (i.e. number of trials back from current trial). It shows that choices were biased away from the sides associated with previously presented stimuli. This repulsive side bias captured the effect of having heard the previous exact stimuli, above an beyond the bias introduced by the category of those stimuli which was captured by the lateral bias (see Methods). It was consistent with an after-effect caused by sensory adaptation in which a strong acoustic power at a given frequency would reduce the likelihood to perceive that frequency in subsequent trials. **c-d**, Fixed side bias (c) and Left and Right lapse rates (d) obtained in GLM fitting for trials following an error (black) or correct response (orange). **e,** Normalized variance of $\gamma^X_t$ (X= S, A, L and T) averaged over animals (see Supplementary Methods) quantifies the overall relative impact of each module (current stimulus, sensory after-effect, lateral bias and transition bias) on the rats choices. **f,** Transfer coefficient $\gamma^T_t \rightarrow \gamma^T_{t+k}$ versus trial lag $k$, quantifying the degree to which the lateral bias at trial $t$ affects the lateral bias on subsequent trials, is calculated separately depending on the outcome of each trial (colored lines show rewarded choices and black lines error choices; see Supplementary Methods for details). Compare with transfer coefficient of the transition bias shown in Fig. 5c. **g**, Transfer coefficient $\gamma^L_t \rightarrow \gamma^L_{t+k}$ for transition bias versus trial lag $k$ for individual animals when trial $t$ was incorrect and trial $t+1$ was correct. The coefficient at $t$+2 was significantly larger than zero for all $n$ = 10 rats (Wald test $p < 0.003$). **h**, Average coefficients for transitions $T^{++}$ ($n$ = 10 animals) obtained when the GLM is fitted separately depending on the outcome of last two trials (see inset; e.g. −+ represents that *t-2* was incorrect and *t-1* was correct). When the last two trials were correct (++), the $T^{++}$ occurring in the last ten trials had a strong influence (dark orange), whereas when the last trial was an error (+− or −−) they had no influence (gray and black dots). Crucially, the influence of previous $T^{++}$ transitions was again strong, when the choice followed a −+ sequence (an error followed by a correct response; light orange dots), confirming the ability of the animals to rapidly recover the transition bias lost after the error, when the subsequent trial was rewarded. Error bars in all panels indicate the first and third quartiles.

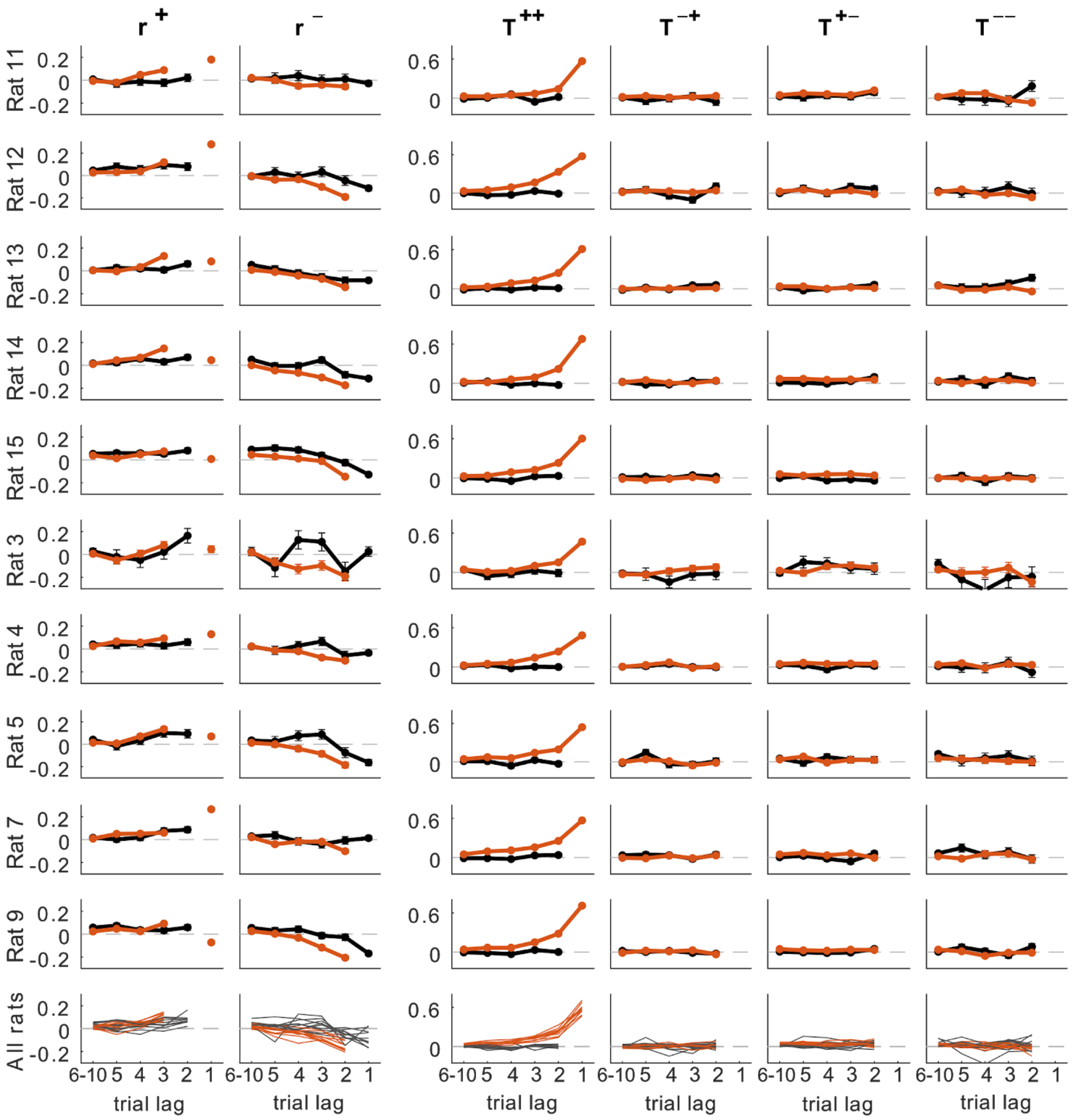


**Supplementary Figure 6. GLM results for individual animals.** GLM weights obtained from each animal in Group 1 (rows) when separately fitting the after-correct (orange) and after-error choices (black). **First two columns:** Lateral weights from previously rewarded ($r^+$) and unrewarded ($r^-$) trials. **Last four columns:** Transition weights computed separately for $T^{++}$ (rewarded-rewarded), $T^{-+}$ (error-rewarded), $T^{+-}$ (rewarded-error) and $T^{--}$ (error-error).

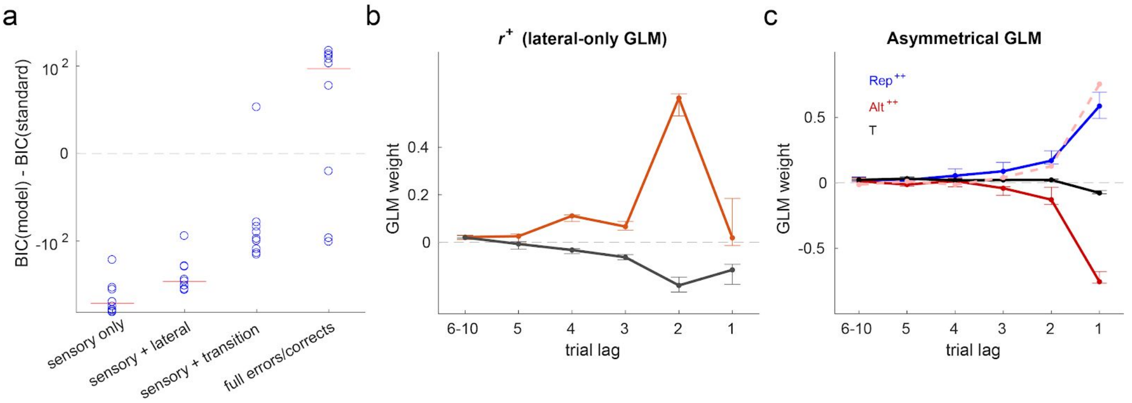


**Supplementary Figure 7. GLM of rat perceptual history bias. a,** Model comparison between the full GLM model and reduced versions of the model with only sensory module (*sensory only*, using only sensory information from current trial), with sensory and lateral modules (*sensory* + *lateral*), with sensory and transition modules (*sensory* + *transition*), and the full GLM fitted separately for trials following correct responses and errors (*full error/corrects*). Comparison is quantified by the difference between Bayesian Information Criterion (BIC) with respect to the full model. Points represent individual animals and vertical lines indicate mean over rats. **b**, Decision weights for lateral regressors $r^+$ (black) and $r^-$ (gray) in the reduced sensory + lateral GLM fitted for all trials. The strong peak at *t-2* for post-correct trials is incompatible with a decaying attraction or repulsion effect. The peak is caused by the large $T^{++}$ weight at *t-1* obtained in after-correct trials in the full model (Fig. 4b left). As explained in the Supp. Methods, regressors $T^{++}_{t-1}$ and $r^{+}_{t-2}$ are identical when computed only in post-correct trials. **c,** GLM weights for a model that used separate regressors for correct repetitions ($Rep^{++}$, blue) and correct alternations ($Alt^{++}$, red solid curve). A third generic transition regressor (*T*, black) was also added. Amplitude for $Rep^{++}$ and $Alt^{++}$ weights were comparable (dashed red shows inverted $Alt^{++}$ weights), showing that correct repetitions and alternations generate symmetrical biases, supporting the idea of single transition bias. The standard GLM with a single transition bias provided a better account of rat decisions than the modified model with separate $Alt^{++}$ and $Rep^{++}$ regressors (*BIC* difference > 20 for all animals, when fitting both models separately on trials following correct responses and errors).

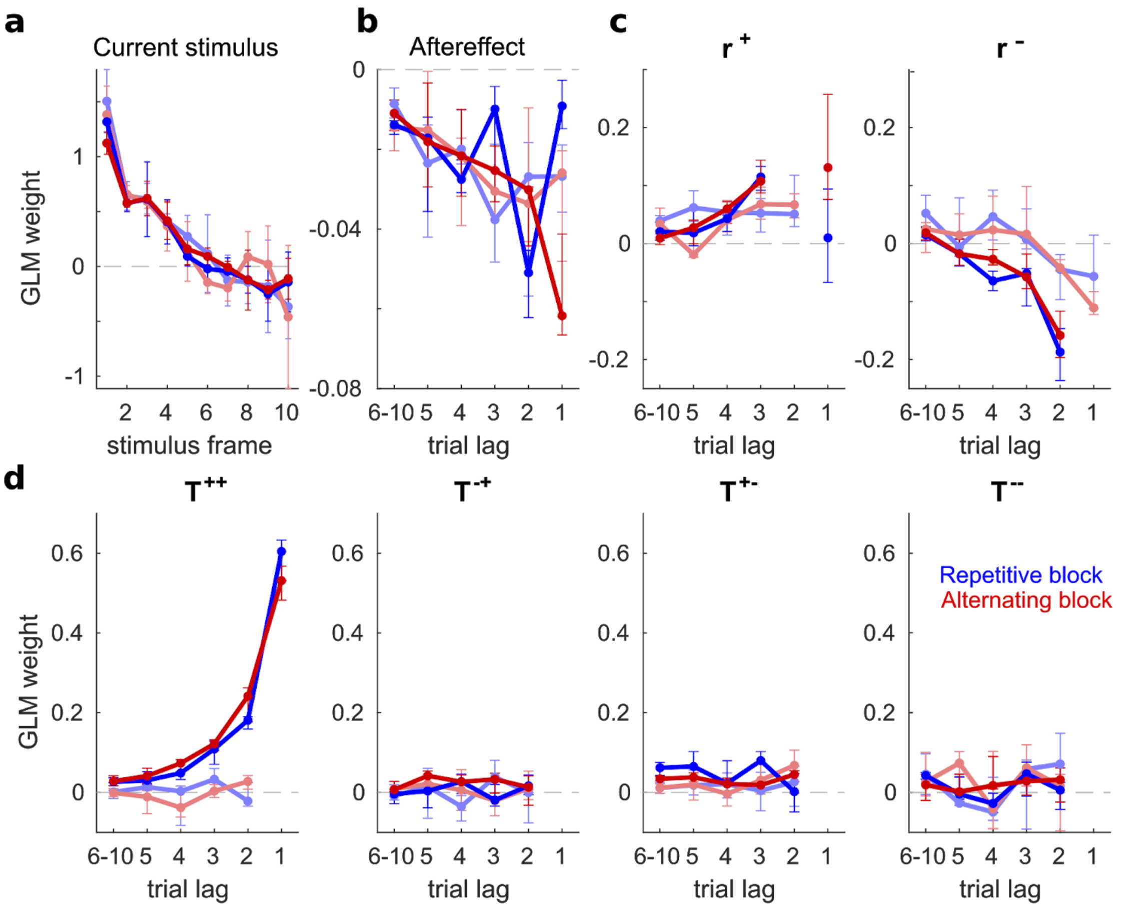


**Supplementary Figure 8. Rats use the same strategy in the two blocks.** Average coefficients obtained in the GLM when separately fitting trials in the Repeating block (blue lines) and in the Alternating block (red lines). As in Fig. 4, weights were fitted separately for after-correct trials (bright colors) and after-error trials (light colors). **a,** Frame-by-frame influence of the current stimulus onto rats decisions. **b,** Influence of the sensory after-effect caused by the net sensory evidence of previous trials as a function of trial lag (i.e. number of trials back from current trial). **c,** Influence of the response side (Left vs Right) from previously rewarded ($r^+$, left panel) and unrewarded ($r^-$, right panel) trials. **d,** Influence of previous transitions (repetition vs. alternation) computed separately for $T^{++}$ (a rewarded trial followed by a rewarded trial), $T^{-+}$ (error-rewarded), $T^{+-}$ (rewarded-error) and $T^{--}$ (error-error). Points in all panels show median coefficients across animals (Group 1, $n$ = 10) and error bars indicate first and third quartiles.

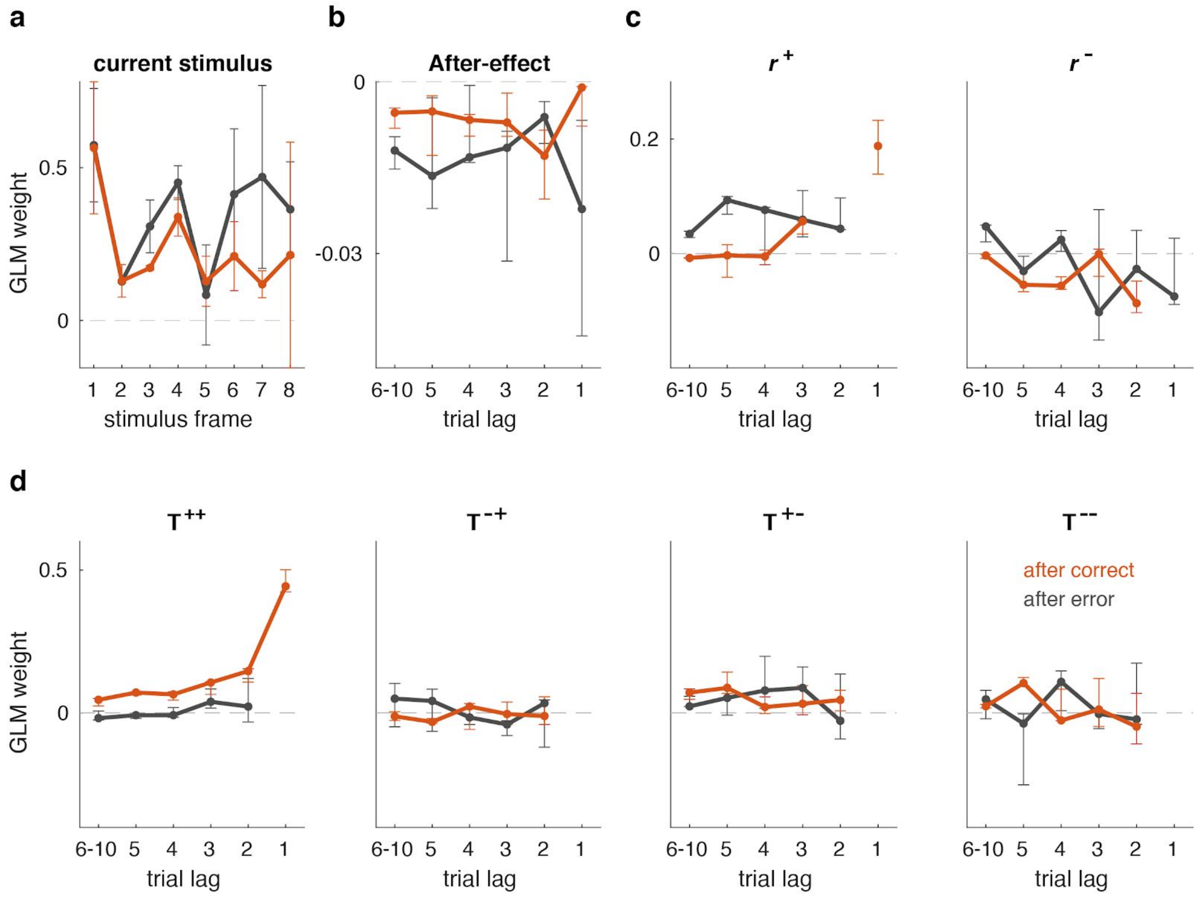


**Supplementary Figure 9. GLM for rats performing an acoustic intensity discrimination task.** Same as Supplementary Fig.8 but for animals in Group 2 (*n* = 6 animals) running the Interaural Level Difference discrimination Task. Average coefficients obtained in the GLM when separately fitting the choices in trials after a correct (orange) and error response (black). Points in all panels show median coefficients across animals and error bars indicate first and third quartiles.

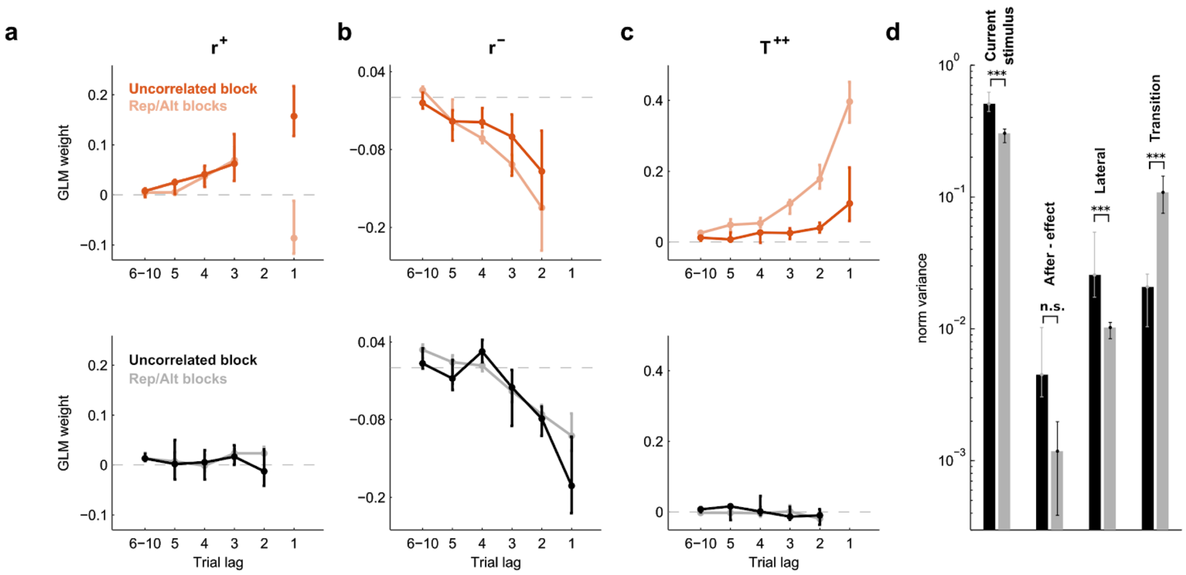


**Supplementary Figure 10. History effects in uncorrelated stimulus sequences. a-c,** Average GLM weights (Group 3, *n* = 9 animals) during initial sessions with uncorrelated stimulus sequences (light colored lines) and during subsequent correlated sessions with Repetitive and Alternating blocks (dark colored lines). Choices were fitted separately after-correct (top row) and after-error trials (bottom row). The lateral weights **(a-b)** do not vary substantially after introducing correlated sequences (except the coefficient of $r^{+}_{t\text{-}1}$). In contrast, the magnitude of the weights of the $T^{++}$ transitions **(c)** after correct increased substantially, whereas the magnitude after errors remained at zero. Error bars show the 1st and 3rd quartiles. **d**, Normalized variance of $\gamma^{X}_{t}$ (X= S, A, L and T) averaged over animals of each module (see labels over bars) before (black) and after introducing sequence correlations (gray). Error bars indicate the 1st and 3rd quartiles.

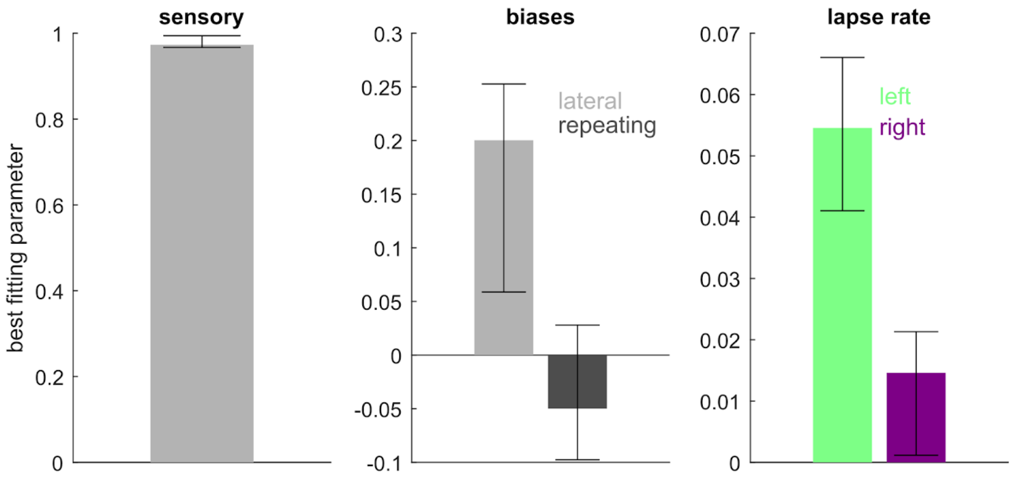


**Supplementary Figure 11. Dynamical model parameters.** Best fitting parameters for sensory sensitivity ($\mu$, left panel), fixed lateral bias and fixed repeating bias ($\beta^{L}$ and $\beta^{T}$, center panel) and lapse rates (right panel). Bars in all panels show median coefficients across animals (Group 1, *n* = 10 animals) and error bars indicate first and third quartiles.

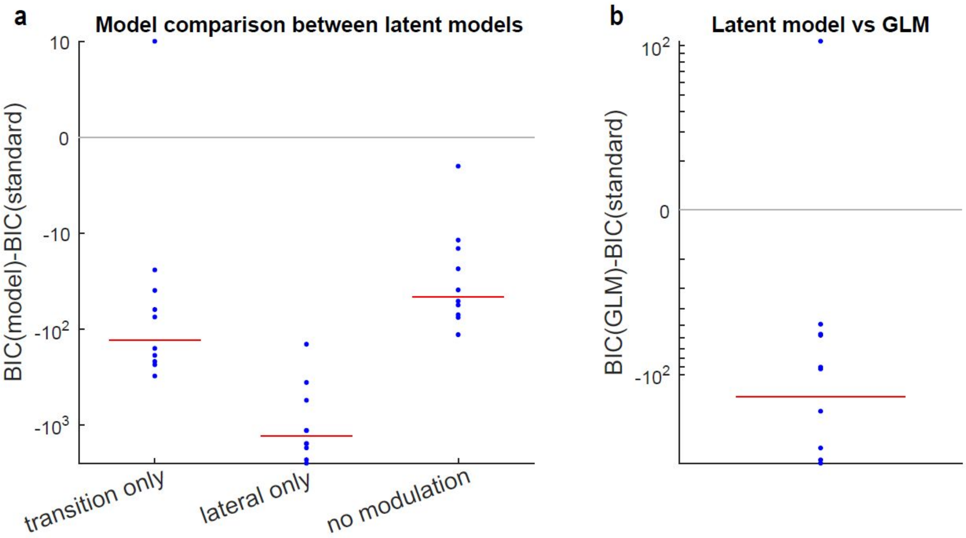


**Supplementary Figure 12. Comparison of dynamical model. a,** the Standard dynamical model of rat behaviour (latent variables: $z^L$, $z^T$, $c^T$) described in Figure 6 is compared to model variants: *transition only* (model without lateral model, i.e. with only $z^T$ and $c^T$), *lateral only* (model without transition bias, i.e. with $z^L$ only) and *no modulation* (both transition and lateral module but no modulation signal, i.e. $z^L$ and $z^T$). Blue points indicate the difference between Bayesian Information Criterion (BIC) of the plotted model and the Standard model for each individual rat (Group1, $n$=10 animals). Red vertical lines indicate rat average. Large negative values indicate very strong evidence in favor of Standard model. **b,** the standard dynamical model is compared with GLM fitted separately for trials after-correct and after-incorrect. The GLM is heavily penalized for the large number of parameters. All rats but one show strong evidence in favor of the Standard dynamical model.

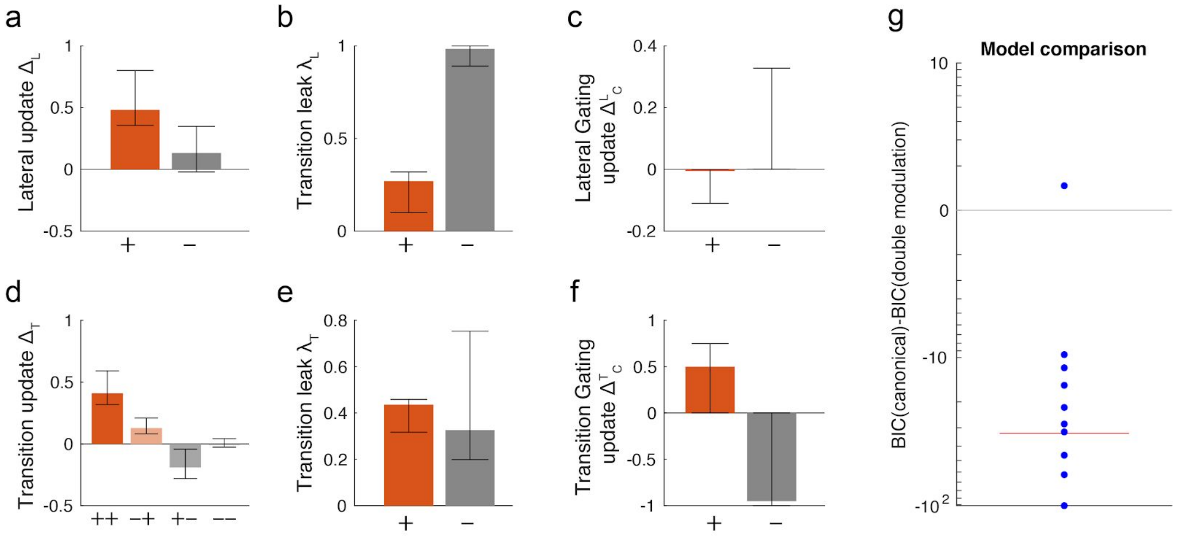


**Supplementary Figure 13. Dynamical model with outcome-dependent gating in both lateral and transition bias.** The model differs from the standard dynamical model by adding the same outcome-dependent gating mechanism to the lateral evidence $z^L$. Best-fitting values of the update parameters in the generative model (median across rats). **a,** lateral evidence update $\Delta_L$ depending on trial outcome, i.e. correct (+) vs. error (−). **b,** outcome-dependent leak of the lateral bias $\lambda_L$. **c,** outcome-dependent update of the transition gating signal $\mathbf{\Delta}^L{}_C$ for the lateral evidence. This extra parameter shows null or close to null gating of the lateral module for most of the rats, both after correct and after error trials, suggesting that this modulatory effect provided little if any benefit to data-fitting. **d,** transition evidence update $\Delta_T$ depending on outcome of the last two trials (++,−+,+−,−−). **e,** outcome-dependent leak of the transition bias $\lambda_T$. **f,** outcome-dependent update of the transition gating signal $\mathbf{\Delta}^T{}_C$ of the transition evidence. A value of -1 correspond to an extinction of the gating signal on the subsequent trial (i.e. a full blockade of the corresponding bias), while +1 correspond to full recovery of the bias (i.e. gating equal to its maximum value of 1). **g,** model comparison shows a minor improvement of the double-gating model compared to Standard model. Values for the rest of parameters are consistent with results from the Standard model. Bars show median coefficients across animals (Group 1, $n$=10 animals) and error bars indicate first and third quartiles.

### Supplementary methods

#### 1 Behavioral training procedure

Training started with three sessions of handling and habituation to the behavioral box followed by five training stages.

- First stage: A lateralized sound was played from one speaker. 200ms after stimulus onset, water was delivered from the corresponding side port independently of the animals' behavior. When the animals collected the reward, the sound was stopped and the trial finished. The side of the sound and water delivery alternated in blocks of 50 trials. After 400 trials the side was randomly interleaved. The first session lasted one night (with free access to food). Animals advanced to the next stage after completing 600 trials across sessions.

- Second stage: Rats learned to self-initiate the trials by poking in the center port. An LED in the center port was lit after the inter-trial-interval ($\sim$1 s) indicating the animal they could start a new trial. The LED remained on for 300ms cueing the animals a fixation period in which they had to remain inside the port. Just after the LED switched off, the stimulus was presented (i) in both speakers in the frequency discrimination task and (ii) in the corresponding speaker in the level discrimination task. The rat withdrew from the center port at any time after stimulus onset, but the sound went on until the animal entered in a side port. Poking in the correct side port resulted in rewarded delivery, whereas incorrect poking was punished with a time out 5s and with a bright light. When the rats had learned the task (performance higher than 90%) we moved to the third stage.

- Third stage: The sound now stopped when the animals left the center port simultaneously the center led offset. The rewards were only available for 4s. When 85% performance was reached animals, advanced to the next stage.

- Fourth stage: the stimulus difficulty was slightly increased on each session by presenting intermediate stimulus evidences. The final evidence values were adjusted to sample the psychometric curve uniformly. Once the final values were achieved the rats went on to the last stage.

- Fifth stage: the rats were exposed to the Repeating and Alternating blocks while keeping the trial structure unchanged.

In animal groups 2 and 3, stages three and four were reversed. Moreover, the decrease in the duration of the stimulus, from lasting until the animal poked on the side ports to lasting only until the withdrawal from the center poke, was done progressively across and within sessions using an automated adaptive method. The complete training process lasted on average 59 sessions per animal.

#### 2 GLM analysis of influence of previous history

##### 2.1 GLM model

In order to understand the relative influence of the different features of recent history and current sensory stimulation onto rats decisions, we built a Generalized Linear Model (GLM) where these different features are linearly summed to give rise to the probability that the rat selects the Right port in each trial $t$ (Eq. 1) [Corrado et al., 2009, Busse et al., 2011, Gold et al., 2008, Akrami et al., 2018, Braun et al., 2018, Fründ et al., 2014, Nogueira et al., 2017] Feature weights were fitted to individual rat decisions. Features were grouped in different modules:

1. The **Sensory module** contains the dependencies on the current and previous acoustic stimuli.

   (a) **Current Stimulus terms** summarize the influence of the *current* auditory stimulus on the decision, i.e. the strength of stimulus-response associations. The module includes a regressor $S_{t,f}$ representing the $t$-th trial intensity difference between the two sounds in frame $f$, for $f = 1, 2 \dots 8$ (the influence of frames with $f > 8$ could be neglected as reaction times longer than 400 ms, i.e. 8 frames, represented on average less than 2.9% of the trials).

   (b) **Stimulus after-effect terms** measure biases due to the presentations of previous stimuli. Features consisted of the overall sensory evidence of each trial $S_t^{sum} = \Sigma_f S_{t,f}$, i.e. the sum of stimulus evidence over all frames, for previous trials $t = -1, -2, \dots -10$. This feature captures any specific potentiation (resp. adaptation) to previously heard tones, that would lead to a bias towards (resp. away from) the side associated with the tone that dominated in previous trials. The regressors capturing this feature were built as for the Lateral module (see below). Although the physical stimulus was strongly correlated with the rewarded side (i.e. stimulus category), fluctuations across frames and variability on the stimulus duration allowed to isolate the effect of the physical stimulus from the category (e.g. for stimuli with low stimulus strength, the sign of the overall physical stimulus $S_t^{sum}$ was often at odds with the stimulus category, facilitating the separate weighting of the two features).

2. The **Lateral module** summarizes the contribution of the side and outcome of responses from previous trials to the lateral bias in each response, i.e. a history-dependent approach or avoidance bias towards either port. The module includes the following regressors:

   (a) The side of correct responses $r_t^+ = r_t \delta_{O_t,1}$, taking value -1 for correct Left responses, 1 for correct Right responses, and 0 for incorrect responses. The output variable $O_t$ equals $+1$ when the trial $t$ was rewarded and -1 when it was not. The $\delta$ is the Kronecker operator, i.e. $\delta_{i,j} = 1$ if $i = j$, and $\delta_{i,j} = 0$ if $i \neq j$. Positive weighting of this feature capture a propensity to opt for previously rewarded responses (i.e *win-stay*).

   (b) The side of incorrect responses $r_t^- = r_t \delta_{O_t,-1}$, taking value -1 for incorrect Left responses, 1 for incorrect Right responses, and 0 for correct responses. Negative weighting of this regressor would capture a propensity to go away from previously non-rewarded responses (i.e *loose-switch*).

   We used regressors $r_t^o$, with $o \in \{+,-\}$, from each of the five preceding trials (i.e. $t = -1, -2, \dots -5$) to measure the influence of previous history over that short window. Another grouped regressor $r_t^o$ was created by summing the corresponding value of trials $t = -6, -7, \dots -10$ .

3. The **Transition module** describes how repetitions and alternations in previous trials affect the propensity to select the same response as in the previous trial (i.e. repeat) or the other one (i.e. alternate). Transition regressors, taking values $+1$ for repetition and -1 for alternation, were separated depending on the outcome on the two successive trials making up the transition:

   (a) The transitions between two consecutively correct responses $T_t^{++} = r_{t-1}^+ r_t^+$, taking value $+1$ for repetition between the two correct trials (denoted in Figs. 2 and 7 as $X^+X^+$ and meaning two consecutive correct Left responses or two consecutive correct Right responses), -1 for alternation between the trials ($Y^+X^+$, Left followed by Right or Right followed by Left, both correct), and 0 if either trial $t$ or $t-1$ was incorrect.

   (b) The transitions from a correct to an incorrect response $T_t^{+-} = r_{t-1}^+ r_t^-$, taking value $+1$ for repetition between the two trials (e.g. Left correct followed by Left incorrect), -1 for alternation between the trials (e.g. Left correct followed by Right incorrect), and 0 for other trials.

   (c) The transitions from an incorrect to a correct response $T_t^{-+} = r_{t-1}^- r_t^+$, taking value $+1$ for repetition between the two trials (e.g. Left incorrect followed by Left correct), -1 for alternation between the trials (e.g. Left incorrect followed by Right correct), and 0 for other trials.

(d) The transition between two consecutively incorrect responses $T_t^{--} = r_{t-1}^- r_t^-$, taking value $+1$ for repetition between the two incorrect trials, -1 for alternation between two incorrect trials, 0 if either trial $t$ or $t-1$ was correct.

As for the lateral module, we used one regressor for each of the five preceding trials and a grouped regressor for the summed value of trials $t = -6, -7, \ldots - 10$, obtaining a total of $4 \times 6 = 24$ regressors $T_t^{o,q}$, with $o, q \in \{+, -\}$. Since these regressors implement a tendency to repeat the previous response or alternate based on recent history, we multiplied each regressor by the previous response $r_{t-1}$ to obtain the transition bias $\gamma^T$, i.e. the tendency to go Left or Right in the subsequent trial (see Eq. 1).

Note that regressors $r_{t-2}^+$ and $r_{t-2}^-$ were excluded from the fitting because they are redundant with transition regressors for the immediately preceding trial (see section '*Indeterminacy on the Influence of transition at previous trial vs side of response two trials back*').

The probability of a Right response at trial $t$ was then modeled as a sigmoidal function of the weighted sum of all regressors from all modules:

$$\begin{cases} p(r_t = +1|\boldsymbol{\omega}, \boldsymbol{\pi}, \beta) = \pi_R + (1 - \pi_L - \pi_R)\ \Phi(y_t) \\ y_t = \sum_f \omega_f^S S_{t,f} + \sum_k \omega_k^A S_{t-k}^{sum} + \sum_{k,o} \omega_{k,o}^L r_{t-k}^o + (\sum_{k,o,q} \omega_{k,o,q}^T T_{t-k}^{o,q})\, r_{t-1} + \beta \end{cases} \quad (1)$$

$\pi_L$ and $\pi_R$ represent the lapse rates for Left and Right responses (i.e. a fixed probability to choose the Left or Right response at any trial independently of current sensory evidence and previous history); $\Phi$ is the cumulative of the standard normal function; $\omega_f^S$, $\omega_k^A$, $\omega_{k,o}^L$ and $\omega_{k,o,q}^T$ represent the weights of the regressors associated to the current stimulus, previous stimulus after-effect, lateral and transition biases, respectively ($o, q \in \{+, -\}$). $\beta$ is the fixed side bias representing the preference of the animal, independent of recent history, to choose one response over the other. Weights $\boldsymbol{\omega}$, overall bias $\beta$ and lapse rates $\boldsymbol{\pi}$ were fitted to the decisions of each rat individually, using all valid trials across sesssions. We also fitted weights separately for trials following correct responses and for trials following errors. The fitting procedure involves a generalized Expectation-Maximization algorithm, implemented in Matlab [Fründ et al., 2014]. We used L2 regularization with a regularization term $\lambda = 1$.

#### 2.2 Impact of each module's estimated bias

In order to assess the relative impact of each of the modules onto the decision we calculated the trial-to-trial estimate of the contribution of each module's bias using the estimated weights $\hat{\omega}$ (i.e. fitted weights) to each regressor and summing over all regressors:

$$\begin{array}{lcll} \gamma_t^S & = & \sum_f \hat{\omega}_f^S S_{t,f} & \text{(current stimulus evidence)} \\ \gamma_t^A & = & \sum_k \hat{\omega}_k^A S_{t-k}^{sum} & \text{(previous stimulus after-effect)} \\ \gamma_t^L & = & \sum_{k,o} \hat{\omega}_{k,o}^L r_{t-k}^o & \text{(lateral bias)} \\ \gamma_t^T & = & \sum_{k,o,q} \hat{\omega}_{k,o,q}^T T_{t-k}^{o,q} r_{t-1} & \text{(transition bias)} \end{array} \quad (2)$$

We then computed the variance of each of the biases $\gamma_t^X$ ($X = S, A, L$ and $T$) and normalized by the total variance, i.e. including the explained and unexplained behavioral variability [Fründ et al., 2014]. Indeed, when using the probit link function ($\Phi^{-1}$), the unexplained variability has normal distribution of unit variance; this is because the probit regression model $p(r) = \Phi(\sum_k w_k X_k)$ is equivalent to $r = H(\sum_k w_k X_k + \eta)$, where $\eta$ is some noise process emitted with standard normal distribution.

$$NV_X = \frac{\mathrm{Var}(\gamma^X)}{1 + \sum_{Y \in \{S,A,L,T\}} \mathrm{Var}(\gamma^Y)} \quad (3)$$

Normalized variance $NV_X$ for module $X$ has the great advantage that it is largely invariant to the addition or subtraction of uncorrelated regressors in the GLM, just as the percentage of variance explained in linear regression.

#### 2.3 Indeterminacy on the Influence of transition at previous trial vs side of response two trials back

While the GLM analysis allows to tear apart the effect of lateral and transition history-dependent biases, one indeterminacy remains in this analysis about the respective contribution of the previous transition ($T_{t-1}$, i.e. the transition at $t-1$) and the side of response two trials back ($r_{t-2}$, i.e. the side at $t-2$) when we perform the analysis separately for trials following an error and following a rewarded response. This can be seen in the following way: a weight $\omega$ for $T_{t-1}$ means that if trial $t-1$ is a repetition trial, i.e. $XX$, there will be a bias $w$ towards repeating $X$ at trial $t$. If on the contrary, trial $t-1$ is an alternation trial, i.e. $XY$, there will be a bias $-\omega$ towards repeating $Y$, which is equivalent to a bias $\omega$ towards selecting $X$ at trial $t$ (denoting $X$ the side of response at trial $t-2$ and $Y$ the alternate response). In both cases increasing the weight for $T_{t-1}$ is equivalent to increasing the weight for $r_{t-2}$ of the same amount, i.e. increasing the lateral bias towards the side selected two trials back. When performing the analysis separately for trials after rewarded responses (first four rows in Table 1) and trials after error responses (last four rows in Table 1) there is an equivalence between the following pairs of regressors: after correct responses, regressors $T^{++}_{t-1}$ vs. $r^{+}_{t-2}$ (yellow cells in Table 1), and $T^{-+}_{t-1}$ vs. $r^{-}_{t-2}$ (gray cells); after error responses, regressors $T^{+-}_{t-1}$ vs. $r^{+}_{t-2}$ (blue cells), and $T^{--}_{t-1}$ vs. $r^{-}_{t-2}$ (green cells).

| | | Regressors | | | | | |
|---|---|---|---|---|---|---|---|
| | | $r^{+}_{t-2}$ | $r^{-}_{t-2}$ | $T^{++}_{t-1}$ | $T^{+-}_{t-1}$ | $T^{-+}_{t-1}$ | $T^{--}_{t-1}$ |
| After Correct | ...$X^+X^+$ | 1 | 0 | 1 | 0 | 0 | 0 |
| | ...$X^-X^+$ | 0 | 1 | 0 | 0 | 1 | 0 |
| | ...$Y^+X^+$ | -1 | 0 | -1 | 0 | 0 | 0 |
| | ...$Y^-X^+$ | 0 | -1 | 0 | 0 | -1 | 0 |
| After Error | ...$X^+X^-$ | 1 | 0 | 0 | 1 | 0 | 0 |
| | ...$X^-X^-$ | 0 | 1 | 0 | 0 | 0 | 1 |
| | ...$Y^+X^-$ | -1 | 0 | 0 | -1 | 0 | 0 |
| | ...$Y^-X^-$ | 0 | -1 | 0 | 0 | 0 | -1 |

Table 1. Correspondence between $T_{t-1}$ and $r_{t-2}$ regressors

In the GLM analysis, we thus computed a single weight for each pair of identical regressors (e.g. $T^{++}_{t-1}$ and $r^{+}_{t-2}$ after correct trials) that accounts for the contribution of both regressors in the pair to the history-dependent bias. We then attributed a posteriori the fitted weight to either the corresponding transition regressor or lateral regressor, selecting the one that was most compatible with its value in the preceding trial (i.e. $r_{t-3}$ for $r_{t-2}$ ; $T_{t-2}$ for $T_{t-1}$ ). In all four cases except $T^{++}_{t-1}$, the lateral regressor was selected, as it provided a nice interpolation of the corresponding values from trials $t-1$ and $t-3$, whereas the corresponding transition regressor ($T^{-+}$, $T^{+-}$, and $T^{--}$) had values non-significantly different from zero for earlier trials (see lighter dots in Figure 4a). By contrast, the transition regressor was selected in the case of $T^{++}_{t-1}$, as it corresponded approximately to an exponential extrapolation of the weights of $T^{++}$ for earlier trials (see lighter dot in Figure 4b left), and was not compatible with the much smaller values of $r^{+}_{t-3}$. Moreover, in the *sensory+lateral* GLM model that did not feature a transition module, the fitted kernels for the lateral module were non-monotonic, because the weight for $r^{+}_{t-2}$ was much larger than for $r^{+}_{t-1}$ (Supp. Fig. 7b). This peculiar effect of a stronger impact of an event further in time is readily accounted for by the fact that peak weight mostly represents the influence of the previous transition, rather than of the side of response two trials back.
The weights attribution was supported by correlation analyses with the neighbouring weights of $T_{t-2}$ and $r_{t-3}$ across the 25 animals. After correct trials, the weight of the undetermined regressor

($T^{++}_{t-1}$, $r^{+}_{t-2}$) correlated strongly with $T^{++}_{t-2}$ ($r = 0.67$, $p < 0.001$) but not with $r^{+}_{t-3}$ ($p > 0.1$). By contrast, the weight of ($T^{-+}_{t-1}$,$r^{-}_{t-2}$) correlated with $r^{-}_{t-3}$ ($r = 0.60$, $p = 0.0016$) but not with $T^{-+}_{t-2}$ ($p > 0.2$). After error responses, the weight of ($T^{+-}_{t-1}$,$r^{+}_{t-2}$) correlated with $r^{+}_{t-3}$ (r=0.71, p<0.001) but not with $T^{+-}_{t-2}$ ($p > 0.2$). The weight of ($T^{--}_{t-1}$, $r^{-}_{t-2}$ ) did not correlate significantly with either $T^{--}_{t-2}$ or $r^{-}_{t-3}$ ($p > 0.2$).

#### 2.4 Alternative GLM models

As control models, we also fitted rat individual data to the following variants of the GLM: (1) a *sensory* model in which all regressors related to history (after-effect, lateral and transition) were removed, (2) a *sensory+lateral* model in which all history regressors except the lateral were removed and (3) a *sensory+transition* all history terms except the transition were removed. For both *sensory* and *sensory+transition* models, we added as a regressor the response at the previous trial $r_{t-1}$, in order to grasp any overall fixed repeating bias. Model comparison was performed using Bayesian Information Criterion (BIC) (Supplementary Fig. 7a). Comparison using the corrected Akaike Information Criterion (AICc) provided similar results.

We also assessed whether both repeating and alternating events played a separate role in the formation of the transition bias, and whether these roles were mirror images of each other suggesting that animals were indeed conceptualizing both patterns, i.e. Rep vs Alt, as the opposite sides of the same bias. For this, we compared our canonical model with a variant model where the transition regressors $T^{o,q}_t$ described above were replaced by the following ones (Supplementary Fig. 7c):

1. The repetition of two consecutively correct responses $Rep^{++}_t$ taking value 1 for repetition between the two correct trials (i.e. $X^+X^+$), and 0 otherwise.
2. The alternation of two consecutively correct responses $Alt^{++}_t$ taking value 1 for alternation between the two correct trials (i.e. $Y^+X^+$), and 0 otherwise.
3. The generic transition between successive responses $T_t$, independently of the outcome of these responses, taking value 1 for any repetition (i.e. $XX$) and -1 for any alternation (i.e. $YX$).

#### 2.5 Testing the complete reset hypothesis versus the gating hypothesis

If after an error at a given trial $t$, the transition bias is *completely reset*, then events occurring before trial $t$ should not be able to generate a transition bias for subsequent trials (Fig. 5a). In order words $\gamma^T_t$, which integrates the transition features up to trial $t-1$, should not influence decisions at $t+1$, +2, and so on. Alternatively, it could be that the bias $\gamma^T_t$ becomes zero after an error because the information about previous transitions is gated off (i.e. not used) but it is not erased (Fig. 5b). In this case the value of the bias before the error $\gamma^T_t$ is predictive of the value of the bias once the gating is back on. To test these two hypothesis, we ran further analyses quantifying the impact of the transition and lateral biases at trial $t$, $\gamma^T_t$ and $\gamma^L_t$, on choices at trial $t+1$ separately for when trial $t$ was correct and it was incorrect (i.e. $O_t = 1$ and $O_t = -1$). Moreover, we also assessed the impact of the biases a trial $t$ on trial $t+2$, separately for the different outcome combinations in trials $t$, $t+1$ (i.e. $++$, $+-$, $-+$ and $--$) and $t+2$ ($+++$, $++-$, $\ldots$). This was done by fitting a new GLM model for the choices in trial $t+1$ using as regressors the different biases $\gamma^X_t$ at time $t$ ($X = S, A, L$ and $T$) in such a way that the fitted weights $\epsilon^X$ represented modulation of these biases:

$$\begin{cases} p(r_{t+1} = +1|\boldsymbol{\epsilon}, O_t) & = & \epsilon^r \hat{\pi}_R + (1 - \epsilon^l \hat{\pi}_l - \epsilon^r \hat{\pi}_R) \;\; \Psi_{t+1} \\ \Psi_{t+1} & = & \Phi(\sum_{j=0}^{2} \epsilon^S_j \gamma^S_{t+j} + \epsilon^A \gamma^A_t + \epsilon^L \gamma^L_t + \epsilon^T \gamma^T_t r_{t-1} r_t + \xi^L_0 r_t + \xi^T_0 r_{t-1} + \epsilon^B \hat{\beta}) \end{cases} \quad (4)$$

For the sensory bias, we included 3 separate regressors for the trials $t$, $t+1$ and $t+2$, (i.e. $\gamma^S_t$, $\gamma^S_{t+1}$ and $\gamma^S_{t+2}$), checking that responses in each trial are principally driven by sensory evidence at the corresponding trial. The additional weights $\epsilon^B$ and $(\epsilon^l, \epsilon^r)$ were used for modulating the fixed bias and lapses, respectively. We also added additional regressors for response at trial $t$ ($r_t$) and $t-1$ ($r_{t-1} = T_t r_t$) to capture the lateral and repeating bias just due to response at trial $t$ (captured in the weights $\xi^L_0$ and $\xi^T_0$, respectively). We fitted the model separately for post-correct trials ($O_t = +1$) yielding the modulated weights $\boldsymbol{\epsilon}_+ = (\epsilon^S_{0,+}, \epsilon^S_{1,+}, \epsilon^S_{2,+}, \epsilon^A_+, \epsilon^L_+, \epsilon^T_+, \epsilon^B_+, \epsilon^l_+, \epsilon^r_+)$, and similarly for post-error trials ($O_t = -1$) yielding the weights $\boldsymbol{\epsilon}_-$.

We then fitted the choices in trial $t+2$ using the same model except that we added as an extra regressor the response at $t+1$:

$$\begin{cases} p(r_{t+2} = +1|\boldsymbol{\epsilon}, O_t, O_{t+1}) = \epsilon^r \hat{\pi}_R + (1 - \epsilon^l \hat{\pi}_l - \epsilon^r \hat{\pi}_R)\ \Psi_{t+2} \\ \Psi_{t+2} = \Phi(\sum_{j=0}^{2} \epsilon_j^S \gamma_{t+j}^S + \epsilon^A \gamma_t^A + \epsilon^L \gamma_t^L + \epsilon^T \gamma_t^T r_{t-1} r_{t+1} + \xi_0^L r_t + \xi_1^L r_{t+1} + \xi_0^T T_t r_{t+1} + \epsilon^B \beta) \end{cases} \tag{5}$$

This fitting was done separately for the the four combinations of the outcome in trials $t$ and $t+1$ yielding four sets of modulating weights $\boldsymbol{\epsilon}_{++}$, $\boldsymbol{\epsilon}_{+-}$, $\boldsymbol{\epsilon}_{-+}$ and $\boldsymbol{\epsilon}_{--}$. Following the same rationale, we also fitted $p(r_{t+3}|\boldsymbol{\epsilon}, O_t, O_{t+1}, O_{t+2})$ and fitted the coefficients separately for the eight combinations of the outcome in trials $t, t+1$ and $t+2$.

As a sanity check, we also computed modulation weights by using biases at trial $t$ to model decisions at trial $t$, which should give $\epsilon^X = 1$ for each weight by construction (by similarity with Eqs. 1 and 2):

$$p(r_t = +1|\boldsymbol{\epsilon}) = \epsilon^r \hat{\pi}_R + (1 - \epsilon^l \hat{\pi}_l - \epsilon^r \hat{\pi}_R)\ \ \Phi(\sum_{j=0}^{2} \epsilon_j^S \gamma_{t+j}^S + \epsilon^A \gamma_t^A + \epsilon^L \gamma_t^L + \epsilon^T \gamma_t^T + \epsilon^B \beta) \tag{6}$$

We defined the *transfer coefficients* $\gamma_t^X \to \gamma_{t+k}^X$ for $k = 0, 1, 2$ and 3 and $X = T$ (Fig. 5c) and $X = L$ (Supp Fig. 5f) as the weights $\epsilon^T$ and $\epsilon^L$ obtained from the fitting of the different trials: trial $t$ coefficients come from Eq. 6, trial $t+1$ coefficients come from Eq. 4, etc. From the four combinations of weights in trial $t+2$ we only showed $\boldsymbol{\epsilon}_{++}$, $\boldsymbol{\epsilon}_{-+}$ and $\boldsymbol{\epsilon}_{--}$ for the sake of clarity. For the same reason only three out of eight combinations of weights in trial $t+3$ were showed: $\boldsymbol{\epsilon}_{+++}$, $\boldsymbol{\epsilon}_{--+}$ and $\boldsymbol{\epsilon}_{---}$. All other sets of weights were consistent with the gating hypothesis.

### 3 Dynamic variable model of behaviour

#### 3.1 Standard model with modulated transition bias

To implement the gating hypothesis in which transition evidence is maintained in memory after errors but transiently does not affect choices, we developed a compact model in which the accumulated transition evidence $z^T$ was passed on from trial to trial depending on whether the last choice was a repetition or an alternation. This variable maintained a running estimate of the transition statistics and its transformation onto the transition bias $\gamma^T$ was gated by second variable $c^T$ by setting $\gamma^T = c^T \times z^T \times r_{t-1}$ . This modulatory variable $c^T$ was updated only based on each trial outcome. Similarly to the transition evidence, the model also contains a variable $z^L$ that maintained the lateral evidence which, in the canonical version of the model (see below), had no modulatory mechanism and its value was therefore simply used as the lateral bias $\gamma^L = z^L$.

Compared with the GLM that required that rats maintained memory of all the features for each of the 10 previous trials in order to make a decision, this latent variable model reduced the working memory load to simply maintaining the value of three latent variables. We used a simple expression for the updating rules of the variables $z^L$ and $z^T$:

$$z_{t+1}^L = (1 - \lambda_L^{O_t}) z_t^L + r_t \Delta_L^{O_t} \tag{7}$$

$$z_{t+1}^T = (1 - \lambda_T^{O_t}) z_t^T + T_t \Delta_T^{O_t O_{t-1}} \tag{8}$$

$\boldsymbol{\lambda}_X = (\lambda_X^+, \lambda_X^-)$, with $X = L, T$, represents the leak parameters which take a different value depending on the outcome of the current trial $O_t$ thus allowing a rapid decay following errors (i.e. an after-error reset could be obtained by $\lambda_X^- = 1$). $\boldsymbol{\Delta}_L = (\Delta_L^+, \Delta_L^-)$ and $\boldsymbol{\Delta}_T = (\Delta_T^{++}, \Delta_T^{+-}, \Delta_T^{-+}, \Delta_T^{--})$ are the update parameters, which take different values depending on the last trial's outcome $O_t$ for the lateral evidence or on the last two trials outcome $(O_t, O_{t-1})$ for the transition evidence. The model assumed the symmetries (1) of the effect of Right ($r_t = +1$) and Left ($r_t = -1$) responses on $z^L$ (Eq. 7) and (2) of the effect of repetitions ($r_t r_{t-1} = +1$) and alternations ($r_t r_{t-1} = -1$) on $z^T$ (Eq. 8).

The modulatory variable $c^T$ was bounded between 0 (no influence of the transition bias onto decision) and 1 (maximal influence onto decision) and followed the following update dynamics:

$$c_{t+1}^T = \begin{cases} c_t^T + \Delta_C^{O_t}(1 - c_t^T) & \text{if } \Delta_C^{O_t} > 0 \\ c_t^T + \Delta_C^{O_t} c_t^T & \text{if } \Delta_C^{O_t} \leq 0 \end{cases} \tag{9}$$

Thus, there were two real-valued parameters $\boldsymbol{\Delta_C} = (\Delta_C^+, \Delta_C^-)$ that determined how the gating variable is updated following a correct trial or an error. Moreover, depending on the value of $\boldsymbol{\Delta_C}$ obtained in the fitting procedure the updating was different: positive $\boldsymbol{\Delta_C}$ values led to an increase in $c^T$, while a negative values led to a decrease (Eq. 9). The value of $\boldsymbol{\Delta_C}$ was bounded between -1 ($c_t^T$ reset to zero) and 1 ($c_t^T$ raised to 1 irrespective of previous value).

The probability of right response was then defined by:

$$p(r_t = +1|z_t^L, z_t^T, \mu, \boldsymbol{\beta}, \pi) = \pi_R + (1 - \pi_L - \pi_R)\ \Phi(\mu\gamma_t^S + z_t^L + \beta_L + (c^T z_t^T + \beta_T)r_{t-1}) \tag{10}$$

For sensory evidence, we used the sensory bias $\gamma_t^S$ computed from equation 2 using weights from the sensory-only GLM model, and the impact of sensory evidence is modulated by parameter $\mu$. The model also included the lateral and repeating fixed bias, $\beta^L$ and $\beta^T$. Initial values for $z^L$ and $z^T$ (for the first trial of a session, or following an invalid trial) were set to 0.

#### 3.2 Fitting procedure

We defined a Gaussian prior with zero-mean diagonal covariance for all model parameters except lapses parameters, and a Dirichlet distribution for the lapse parameters $\boldsymbol{\pi}$:

$$p(\boldsymbol{\Delta}, \boldsymbol{\lambda}, \mu, \boldsymbol{\beta}, \boldsymbol{\pi}) = \mathcal{N}(\boldsymbol{\Delta}, \boldsymbol{\lambda}, \mu, \boldsymbol{\beta}; \boldsymbol{0}, \sigma \boldsymbol{I})\text{Dir}(\pi_L, \pi_R, 1 - \pi_L - \pi_R; \boldsymbol{\alpha^{Dir}}) \tag{11}$$

We used $\sigma = 1$ and $\alpha^{Dir} = (1; 1; 38)$ (yielding an overall prior lapse probability of 5%).

Parameters in the latent model (both with and without modulated biases) were fitted by maximizing the log-posterior of the observed responses of rats (i.e. the sum of the log-prior and the log-likelihood):

$$\ln p\boldsymbol{\Delta}, \boldsymbol{\lambda}, \mu, \boldsymbol{\beta}, \boldsymbol{\pi}|\boldsymbol{R}) = \ln p(\boldsymbol{\Delta}, \boldsymbol{\lambda}, \mu, \boldsymbol{\beta}, \boldsymbol{\pi}) + \sum_t (\delta_{R_t,1} \ln p_t + \delta_{r_t,-1} \ln(1 - p_t)) \tag{12}$$

where $p_t = p(r_t|z_t^L, z_t^T, c_t^L, c_t^T, \mu, \boldsymbol{\beta}, \boldsymbol{\pi})$.

Maximization was performed using function *fmincon* from the Matlab Optimization Toolbox, using a gradient-descent algorithm. An iterative formulation of the gradient over all parameters can be found by an analytical derivation of Eqs. 7-8 and Eq. 12. Invalid trials were excluded from the definition of the log-posterior. To avoid that model capture slow (e.g. session-wise) variations in lateral and repeating bias, quite large in some rats, we enforced that the leak was large enough all by adding the constraint $\boldsymbol{\lambda}_X > 0.1$ for both $X = L, T$ (thus restricting the model to integrate over a window of a few dozens trials at most). We run the minimization procedure with 500 randomly selected initial points.

#### 3.3 Variants of the latent model

We also modeled and compared different variants of the latent model:

1. **Modulated lateral and modulated transition** $[z^L, z^T, c^L, c^T]$**.** We defined a second modulatory signal $c^L$ which followed the same type of dynamics as $c^T$ (Eq. 9) but modulated the lateral evidence $z_L$ (i.e. substitute $z_t^L$ in Eq. 10 by $c_t^L \times z_t^L$). The parameters of of the updating of $c^L$ were fitted independently of the parameters of $c^T$ (see Supplementary Fig. 13).

2. **Modulated lateral only** $[z^L, c^L]$**.** This was a version of the model without transition bias (i.e. no $z^T$ nor $c^T$) but with modulated lateral evidence (see Supplementary Fig. 12).

3. **Modulated transition only** $[z^T, c^T]$**.** This was a version of the model without lateral bias (i.e. no $z^L$ nor $c^L$) but with modulated transition evidence (see Supplementary Fig. 12).

4. **No modulation** $[z^L, z^T]$**.** This was a version of the model with only transition evidence $z^T$ and lateral evidence $z^L$ as latent variables but no modulatory variables $c^T$ nor $c^L$ (see Supplementary Fig. 12).

##### 3.4 Degeneracy considerations

If there is no modulation by gating variables (i.e. if the update parameters $\Delta$ are null, which was the case for at three rats), the system of parameters $\{\boldsymbol{\Delta}, \boldsymbol{\lambda}, \mu, \boldsymbol{\beta}, \boldsymbol{\pi}\}$ becomes redundant because a constant term can be added up to the lateral update parameters and subtracted from the fixed lateral bias leaving the system unchanged. The same is true for the transition update parameters and the fixed transition bias. Formally, the parameter set $\{\boldsymbol{\Delta}_L, \boldsymbol{\lambda}_L, \boldsymbol{\beta_L}\}$ is equivalent to $\{\boldsymbol{\Delta}_L + \boldsymbol{\lambda}_L u, \boldsymbol{\lambda_L}, \boldsymbol{\beta_L} - u\}$ for any real $u$, and the same applies to $\{\boldsymbol{\Delta}_T, \boldsymbol{\lambda}_T, \boldsymbol{\beta_T}\}$. To remove such degeneracy, we imposed the additional constraints $< \Delta_L^{O_t} >_{O_t} = 0$ and $< \Delta_T^{O_t,O_{t-1}} >_{O_t,O_{t-1}} = 0$, i.e that the weighted average value of the lateral update across possible outcomes $O_t$ and of the transition update across combinations of consecutive outcomes $O_t O_{t-1}$ were null.

The constraints on update parameters can be written as:

$$n_+ \Delta_L^+ + n_- \Delta_L^- = 0 \tag{13}$$

$$n_{++} \Delta_T^{++} + n_{-+} \Delta_T^{-+} + n_{+-} \Delta_T^{+-} + n_{--} \Delta_T^{--} = 0 \tag{14}$$

where the prefactor $n_{O_+}$ ( $n_{O_t O_{t-1}}$ ) represents the number of trials yielding each outcome $O_t$ (combination of outcomes $O_t, O_{t-1}$). Using Eqs. 13-14, one parameter $\Delta_x$ in each of the update vectors ($\boldsymbol{\Delta_L}$ and $\boldsymbol{\Delta_T}$) could then be determined by the value of all other parameters. We thus removed it from the list of free parameters and used the following equation to compute the gradient for the other (free) update parameters $\Delta_y$:

$$\nabla_{\Delta^y} . = \frac{\partial .}{\partial \Delta_y} - \frac{n_y}{n_x} \frac{\partial .}{\partial \Delta_x} \tag{15}$$

#### References


[Akrami et al., 2018] Akrami, A., Kopec, C. D., Diamond, M. E., and Brody, C. D. (2018). Posterior parietal cortex represents sensory history and mediates its effects on behaviour. *Nature.*

[Braun et al., 2018] Braun, A., Urai, A. E., and Donner, T. H. (2018). Adaptive History Biases Result from Confidence-weighted Accumulation of Past Choices. *The Journal of neuroscience : the official journal of the Society for Neuroscience*, 38(10):2418–2429.

[Busse et al., 2011] Busse, L., Ayaz, A., Dhruv, N. T. N., Katzner, S., Saleem, A. B., Scholvinck, M. L., Zaharia, A. D., Carandini, M., Schölvinck, M. L., Zaharia, A. D., Carandini, M., and Scholvinck, M. L. (2011). The Detection of Visual Contrast in the Behaving Mouse. *Journal of Neuroscience*, 31(31):11351–11361.

[Corrado et al., 2009] Corrado, G. S., Sugrue, L. P., Brown, J. R., and Newsome, W. T. (2009). The Trouble with Choice: Studying Decision Variables in the Brain. *Neuroeconomics*, pages 463–480.

[Fründ et al., 2014] Fründ, I., Wichmann, F. A., and Macke, J. H. (2014). Quantifying the effect of intertrial dependence on perceptual decisions. *Journal of vision*, 14(7):1–16.

[Gold et al., 2008] Gold, J. I., Law, C.-T., Connolly, P., and Bennur, S. (2008). The Relative Influences of Priors and Sensory Evidence on an Oculomotor Decision Variable During Perceptual Learning. *Journal of Neurophysiology*, 100(5):2653–2668.

[Nogueira et al., 2017] Nogueira, R., Abolafia, J. M., Drugowitsch, J., Balaguer-Ballester, E., Sanchez-Vives, M. V., and Moreno-Bote, R. (2017). Lateral orbitofrontal cortex anticipates choices and integrates prior with current information. *Nature Communications*, 8:14823.
